# MTH1 promotes mitotic progression to avoid oxidative DNA damage in cancer cells

**DOI:** 10.1101/575290

**Authors:** Helge Gad, Oliver Mortusewicz, Sean G Rudd, Ailine Stolz, Nuno Amaral, Lars Brautigham, Linda Pudelko, Kumar Sanjiv, Christina Kaldéren, Ann-Sofie Jemth, Ingrid Almlöf, Torkild Visnes, Niklas Schultz, Johan Boström, José Manuel Calderon Montano, Anna Hagenkort, Petra Groth, Olga Loseva, Camilla Gokturk, Tobias Koolmeister, Prasad Wakchaure, Evert Homan, Cecilia E Ström, Martin Scobie, Holger Bastians, Ulrika Warpman Berglund, Thomas Helleday

## Abstract

**BACKGROUND:** We developed MTH1 inhibitors (MTH1i) TH588 and TH1579 showing broad anti-cancer activity, while structurally distinct MTH1i fail to kill cancer cells. Here, we describe a new role of MTH1 in mitosis and the detailed mechanism of action of TH1579 (karonudib) and other structurally distinct MTH1i.

**MATERIALS AND METHODS:** Cancer cell lines or zebrafish embryos were treated with MTH1i or siRNA targeting MTH1 and analysed primarily by live cell and immunofluorescence microscopy, survival assays, DNA fibre or COMET assays. MTH1 and tubulin interactions were analysed *in vitro* using co-immunoprecipitation and tubulin polymerisation assays.

**RESULTS:** Here, we describe a mitotic role for the MTH1 protein, which binds to tubulin, is required for microtubule polymerisation, correct spindle assembly, mitosis progression and suppression reactive oxygen species (ROS) generation in mitosis. Potent MTH1i display differential abilities to break the MTH1-tubulin interaction and cause mitotic arrest, demonstrating 8-oxodGTPase and mitotic function of MTH1 are mechanistically distinct. TH588 and TH1579 have more profound effect on mitotic arrest than other MTH1i explained by additional direct inhibition of tubulin polymerisation. MTH1i only inhibiting 8-oxodGTPase activity synergize with mitotic poisons.

**CONCLUSIONS:** Efficient MTH1 have a dual mechanism of action: inhibiting mitosis (to generate ROS) and promoting 8-oxodGTP incorporation into DNA during mitotic replication, dependent on ROS generation. Direct inhibition of tubulin polymerisation of TH588 and TH1579 increase their ability to arrest cells and generate ROS in mitosis. Furthermore, non-cytotoxic MTH1 can become effective and increase incorporation of oxidised nucleotides into DNA when combined with sub-therapeutic concentrations of mitotic inhibitors or challenged directly by 8-oxodGTP.

## INTRODUCTION

While targeting DNA repair, or the DNA damage response (DDR), as anti-cancer strategy has been a focus over many years [1], the potential huge benefit it can offer is only now becoming apparent, for instance following the recent results obtained in first-line treatment with the PARP inhibitor (PARPi) olaparib in ovarian cancer [2], especially homologous recombination defective (HRD) cancers [3, 4]. Targeting non-essential DNA repair proteins such as PARP1 is a priority and we and others earlier demonstrated that the non-essential DNA repair protein MTH1 is synthetic lethal with the cancer phenotype and developed potent MTH1 inhibitors (MTH1i) that have potent general anti-cancer activity while being well tolerated [5–8]. Currently, the MTH1i karonudib (TH1579) is evaluated in clinical trials, making MTH1 amongst a very short list of DNA repair targets being evaluated in the clinic [9].

The MTH1 protein is a poorly studied DNA repair enzyme and to date only described to prevent oxidized dNTPs, such as 8-oxodGTP and 2-OHdATP, from being incorporated into DNA [10, 11]. Other functions of MTH1 are unknown. Recently, it was demonstrated that potent MTH1i, structurally distinct from TH588 or TH1579, fail to kill cancer cells; challenging MTH1 as anti-cancer target altogether [12–14]. Here, we address the functions of MTH1 as well as the mechanism of action of TH588 and TH1579 along with other inhibitors to be able to explain the discrepancy between MTH1i. These results are important to increase our understanding on the mechanism of action of MTH1i currently being tested in clinical trials.

## MATERIALS AND METHODS

### Cell culture, Inhibitors, Immunofluorescence, Time-lapse microscopy

Cell lines were cultured at 37°C in 5% CO2 and in media supplemented with 10% FBS and TH588[6], TH1579 [15], TH7238 and AZ cmpd #19, #21, #24 [12] were synthesized in house and prepared as previously reported, while other purchased CENP-E inhibitor (CENP-Ei; GSK923295, SelleckChem), Vincristine sulfate (Sigma Aldrich), Paclitaxel (Sigma Aldrich), Nocadozole (Sigma Aldrich), RO3306 (SelleckChem).

Immunofluorescence microscopy with suspension cells was performed as described previously [16] and the time-lapse was initiated after 30 min after treatment and images of GFP and brightfield channels were acquired every 10 min for 24 h. All other experiments are described in detail in supported materials and methods.

## RESULTS

### MTH1 is required for normal mitotic progression

We and others have previously shown that cancer cells die following MTH1 depletion *in vitro* and *in vivo* [6, 7, 15, 17]. When depleting MTH1 using different siRNA or shRNA *in vitro* or *in vivo* we observed cell division defects indicative of mitotic problems, such as formation of micronuclei or polynucleated cells (Figure 1A-D; Supplementary Figure 1A-J), compared to non-targeting siRNA or shRNA treated cells. When further analyzing mitotic progression by live cell microscopy in H2B-GFP expressing U2OS cells following MTH1 depletion, these cells showed lagging chromosomes (Figure 1E-F, Supplementary Figure 1K). Accurate cell division requires correct establishment of kinetochore-microtubule attachments, therefore, as a readout for correct attachments we next measured the inter-kinetochore distances in sister-chromatids, which was significantly reduced following MTH1 depletion (Figure 1G-H), suggesting MTH1 contributes to the correct establishment of these attachments.

**Fig. 1:**
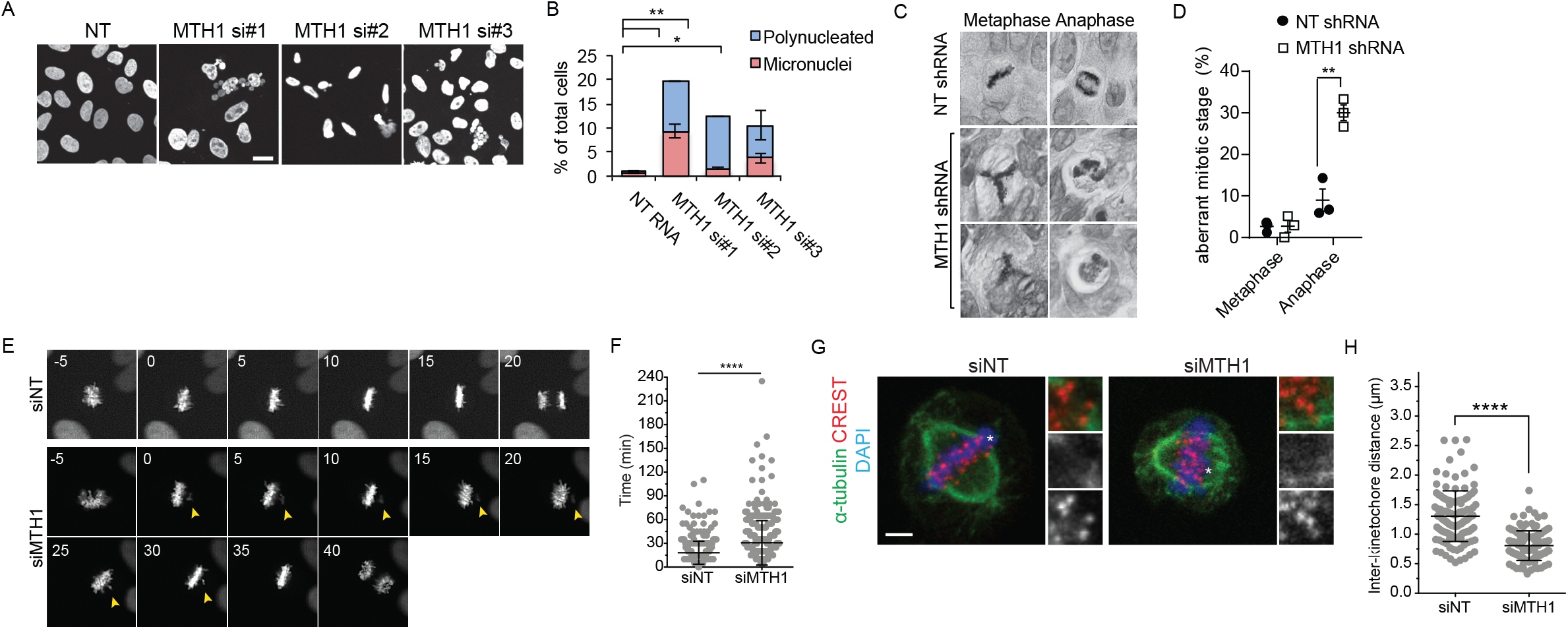
MTH1 knockdown arrest cells, induce polynucleation, perturb mitosis and inhibit chromosome congression in cells. **A,B.** MTH1 knockdown using 3 different siRNAs causes increased fraction of polynucleated cells. U2OS cells were transfected with non-targeting (NT) or MTH1 siRNA for total of 10 days and stained with Histone H3pS10 antibodies and DAPI (A). Scale bar: 20 μm. Quantification of polynuclei and micronuclei following NT or MTH1 siRNA (p<0.05 for polynucleated cells, mean±SD, 2 experiments, >200 cells/sample). **C, D.** SW480 cells with dox-inducible NT or MTH1 shRNA #2 were subcutaneously inoculated in the flank of SCID mice and when the tumour reached approximately 250 mm^3^, doxycyclin was administrated in the drinking water. After additional 5 days, tumours were analysed by immunohistochemistry and cells with aberrant mitotic stages were scored. Data show mean ± SEM of 3 control and 3 MTH1 shRNA tumours. ** p <0.01, Student t-test. **E. F.** Time-lapse of mitosis in siRNA-transfected U2OS cells expressing H2B-GFP. Arrowheads point to lagging or misaligned chromosomes. **E.** Examples of mitotic cells in U2OS cells transfected with either non-targeting (siNT) or MTH1 siRNA. Scale bar: 10 μm. **F.** Time-lapse images of U2OS cells transfected with either non-targeting (siNT) or MTH1 siRNA. Numbers indicate time in minutes relative to metaphase plate establishment (0 minutes). G. Quantification of time to chromosome alignment in the indicated conditions. **H.** Inter-kinetochore distances are reduced upon MTH1 knock-down. Representative figures are shown of the corresponding siRNA-transfected U2OS cells. Examples of kinetochore pairs are shown magnified. **I.** Graph showing inter-kinetochore distance measurements from one representative experiment. Three independent experiments were performed with more than 100 kinetochore pairs measured per experiment. **** p<0.001, Student’s t-test.

In accordance with the MTH1 siRNA results, although much more pronounced, U2OS cells treated with MTH1i TH588 and TH1579 showed an increased time in mitosis shortly after treatment, strongly associated with a delay in chromosome alignment, and their subsequent entry into G1 phase coincided with polynucleation (Figure 2A-C, Supplementary Figure 2A, Movies S1-S3). Interestingly, when we transfected immortalized BJ-hTERT and transformed BJ-RasV12 cells with H2B-GFP and followed their fate in mitosis after TH588 or TH1579 treatment using live cell imaging, we observed similar mitotic phenotypes in transformed BJ-RasV12, but not in BJ-hTERT cells (Figure 2D-E). This is in line with our previous report of increased survival and lower levels of incorporated oxidised nucleotides, such as 8-oxodG, in non-transformed cells [6, 15]. Importantly, as cells were followed continuously by live-cell imaging, the difference between immortalized and Ras-transformed cells cannot simply be explained by Ras-transformed cells having a shorter cell cycle. Furthermore, in accordance with MTH1 siRNA observations, we observed lagging chromosomes (Figure 2B) and a reduced inter-kinetochore distance in comparison to control or paclitaxel treated cells (Figure 2F). Thus, the mitotic phenotype observed after TH588 and TH1579 treatment was similar to the one observed following MTH1 depletion, although much more pronounced.

**Fig. 2:**
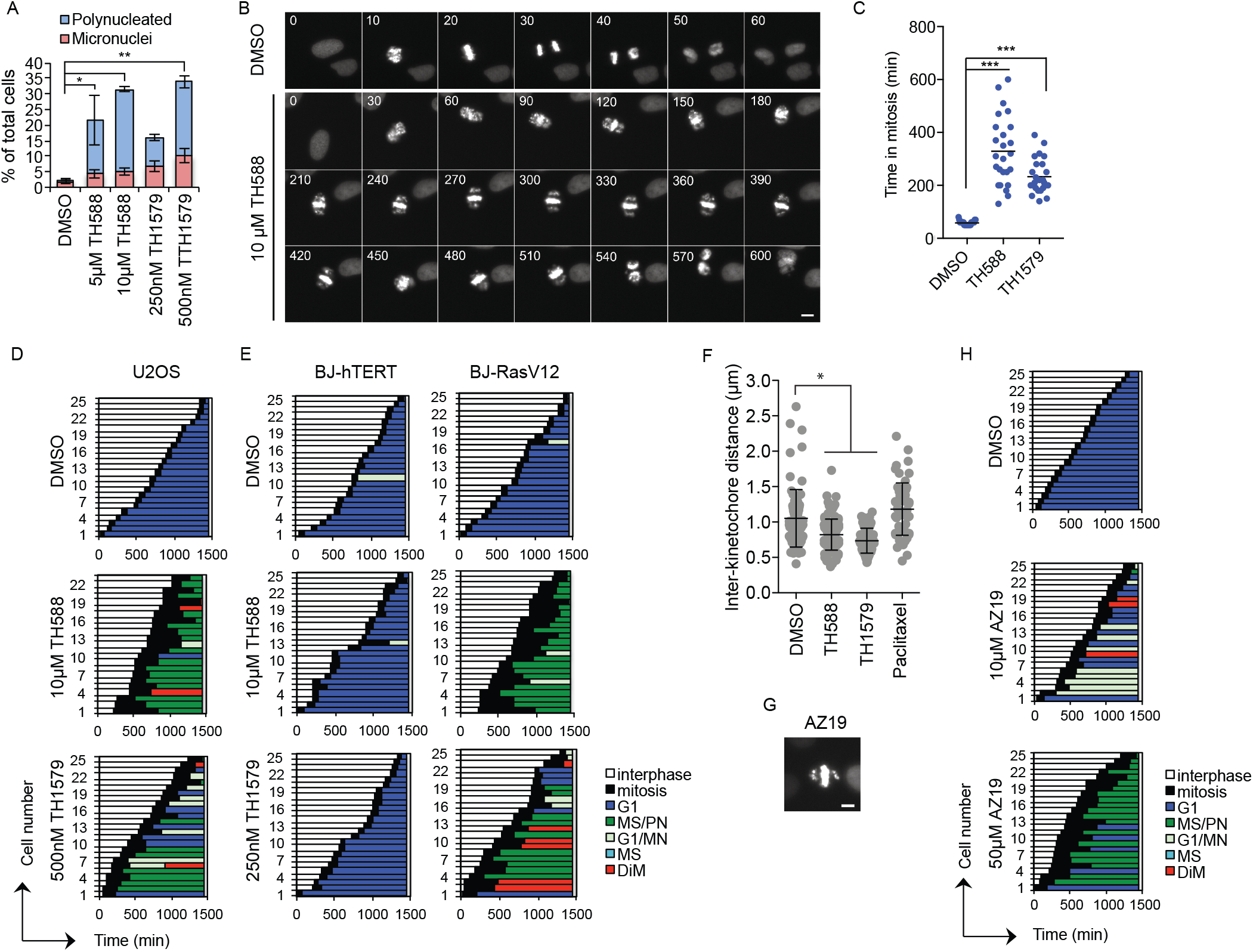
MTH1i TH588 and TH1579 interfere with mitotic progression. **A.** U2OS cells treated with TH588 and TH1579 for 24 h show an increased fraction of polynucleated or micronuclei containing cells. **B-E.** Mitotic progression using time-lapse imaging of U2OS H2B-GFP cells treated with 10 μM TH588 or 0.5 μM TH1579. The time-lapse was initiated 30 min after addition of inhibitors and images were acquired every 10 min for 24 h. Scale bar: 10 μm. **C.** Time in mitosis from pro-phase to end of cytokinesis was quantified in 25 cells per condition. **D.** Individual cells were followed manually and scored for defects in mitosis including mitotic slippage/polynucleation (MS/PN), micronuclei formation (G1/MN), mitotic slippage (MS) or cell death during mitosis (DiM). **E.** Treatment with TH588 and TH1579 cause a mitotic arrest and fail to complete mitosis in Ras-transformed but not immortalized BJ-cells. H2B-GFP expressing BJ-hTERT and BJ-hTERT/SV40T/RASV12 cells were treated with 10 μM TH588 or 250 nM TH1579 and mitotic progression was quantified by time-lapse analysis. Images were acquired every 10 min for 24 h. **F.** Inter-kinetochore distance measurements from one representative experiment. Two independent experiments were performed with more than 50 kinetochore pairs measured per experiment. **G.** High concentration of AZ19 show similar mitotic chromosome congression defect as TH588 and TH1579. U2OS H2B-GFP cells were arrested in mitosis after treatment with 50μM AZ19. Scale bar: 10 μm. **H.** Individual U2OS H2B-GFP cells were followed manually using time-lapse initiated 30 min after addition of 50 μM AZ19 and images were acquired every 10 min for 24 h. Cells were scored for defects in mitosis including mitotic slippage/polynucleation (MS/PN), micronuclei formation (G1/MN), mitotic slippage (MS) or cell death during mitosis (DiM).

Interestingly we observed that a previously reported non-cytotoxic MTH1i, Astra Zeneca compound 19 (AZ19), was cytotoxic at higher concentrations (10-50 μM) and caused a congression defect similar to TH588 or TH1579 treatment or MTH1 depletion, and also resulted in a mitotic arrest and polynucleation (Figure 2G-H; Supplementary Figure 2B-C; Movies S7, S8). This further supports a role for MTH1 in mitotic progression, however given these compounds inhibits the enzymatic function of MTH1 at similar low nM concentrations, this suggests that the role of inhibitors disrupting the mitotic process is distinct from the enzymatic function of MTH1.

### A novel role of MTH1 in microtubule dynamics

Since microtubule dynamics are crucial for accurate mitotic progression [18] and MTH1 depletion interfered with correct kinetochore-microtubule attachments, and caused mitotic delays, lagging chromosomes and polynucleation, we hypothesized that MTH1 plays a role in microtubule dynamics. To test this, microtubule plus end growth rates were determined by tracking EB3-GFP protein in living HCT116 cells. 48h siRNA knockdown of MTH1 did not influence microtubule polymerization rate (Figure 3A), but a prolonged MTH1 siRNA treatment (96 h) of HCT116 cells showed a reduction of the microtubule polymerization rate (Figure 3B, Movie S4). Furthermore, by using fluorescence recovery after photobleaching (FRAP) in U2OS cells expressing mCherry-tagged alpha-tubulin, we could show that tubulin mobility was increased in mitotic cells after MTH1 siRNA depletion which is in agreement with the observed reduction in microtubule polymerization (Figure 3C-E; Supplementary Figure 2D-E). These data suggest a role of MTH1 in tubulin function. Next, we performed pull down experiments using U2OS cells expressing MTH1-GFP or, a catalytically-dead variant of MTH1, GFP-E56A mutated MTH1. Interestingly, alpha-tubulin was found to be bound to MTH1-GFP and significantly less bound to the GFP-E56A mutated MTH1 (Figure 3F-G). The MTH1 and alpha-tubulin interaction was further confirmed using a previously described F3H interaction assay [19, 20]. In brief, a high-affinity GFP binding nanobody (GBP) is coupled to the Lac repressor (LacI) and stably expressed in U2OS F3H cells, which harbor a stably integrated lac operator array (LacO). The GBP-LacI protein binds to the LacO and recruits any GFP-fusion protein that is co-expressed to this LacO region (Supplementary Figure 2F). Expression of MTH1-GFP in these cells resulted in nucleation of alpha-Tubulin filaments spreading from the MTH1-GFP bound LacO region (Figure 3H). This indicates that MTH1 and tubulin interact in cells and that MTH1 potentially can even initiate microtubule nucleation.

**Fig. 3.**
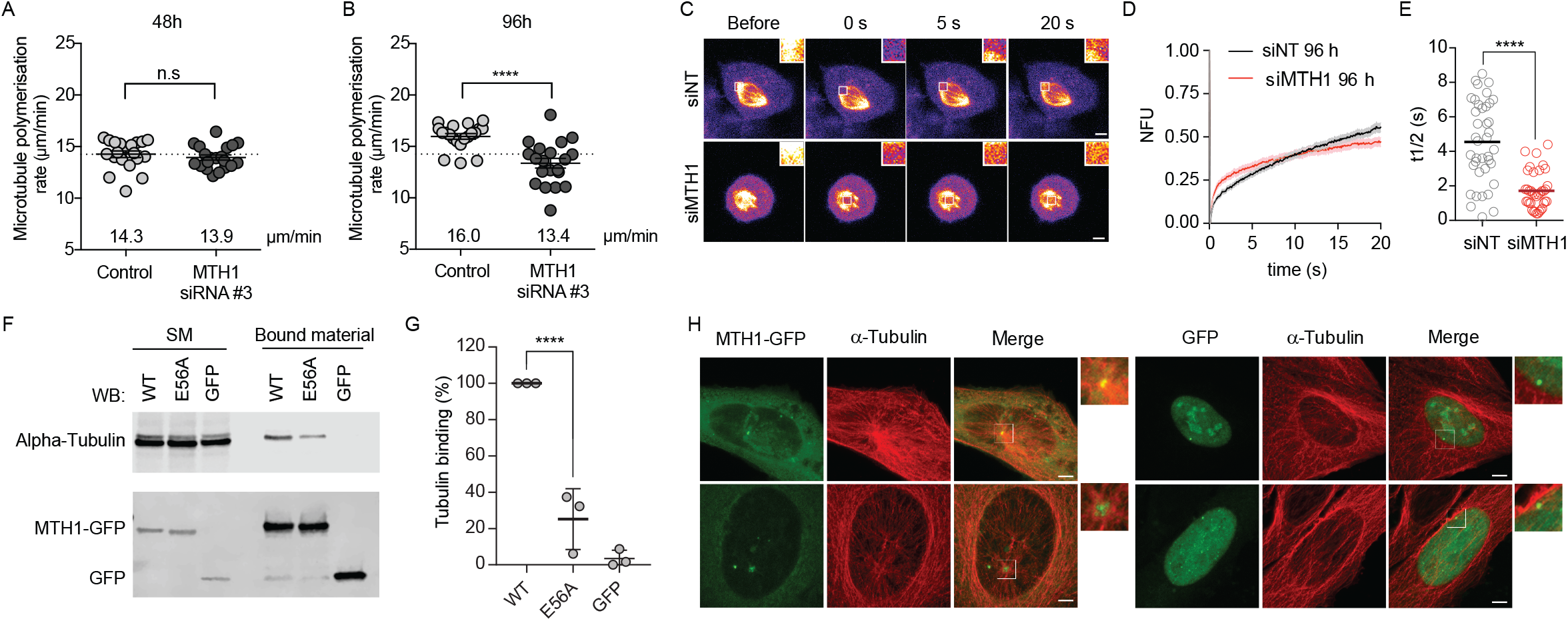
MTH1 knockdown inhibits microtubule dynamics in cells. **A, B.** Long-term knockdown of MTH1 caused decreased MT polymerization rate. Measurement of mitotic microtubule plus end assembly rates in living HCT116 cells, 48 h (A) or 96 h (B) after knockdown of *MTH1* with the indicated siRNAs. Scatter dot plots show average assembly rates (20 individual microtubules per cell, mean +/-SEM, t-test, n=20).). **C.** Representative confocal FRAP image series of mitotic cells transfected with non-targeting (siNT) or MTH1 (siMTH1) siRNA for 96 h. Insets show 3 times magnified bleach area. **D, E.** Quantification of fluorescence recovery after photobleaching in mitotic siNT and siMTH1 cells. **E.** Halftime of recovery (t1/2) was quantified. Scatter dot blots of ≥ 37 cells from 3 independent experiments are shown. Scale bar, 5 μM. (n.s.: non significant, **: p<0.01 ***: p<0.001 ****: p<0.0001, Student’s t-test.). **F, G.** Western blot of immunoprecipitation of MTH1-GFP (WT or E56A mutant) or GFP alone in U2OS cells and probed with anti-Alpha-tubulin and anti-GFP antibodies. Lanes show starting material (SM) and material bound to GFP-trap beads. Bound Alpha-tubulin was quantified from 3 independent experiments (mean±SD, ****: p<0.0001). **H.** Representative confocal images of U2OS F3H cells expressing either GFP or MTH1-GFP, probed with anti α-Tubulin antibody. Inset shows 3 times magnification. Scale bar, 5 μm.

Taken together, these data show that MTH1 interacts with alpha-tubulin and establishes a role for MTH1 in microtubule dynamics, with MTH1 being required for normal mitotic progression.

### TH588 or TH1579 affect tubulin and microtubule dynamics

Previously, TH588 has been reported to inhibit tubulin polymerisation *in vitro* and this was suggested to be responsible for cell killing activity [13]. We can confirm that TH588, and TH1579, inhibit polymerisation of tubulin *in vitro* in a concentration-dependent manner (Fig. 4A-B). Using *in silico* docking studies we identified the colchicine-binding pocket of β-tubulin as the potential binding site of TH588 (Supplementary Figure 3A). In comparison, in silico docking to the colchicine binding pocket suggested TH588 to interact weaker than TH1579 but stronger compared to AZ19 (Figure 4C). To understand how relevant these direct effects on microtubules polymerisation *in vitro* were at therapeutic concentrations in an *in vivo* situation, we subsequently measured microtubule mass and dynamics in cells. Firstly we used a cell-based assay monitoring whole cell microtubule mass and treated HCT116 cells with TH1579, TH588, Paclitaxel, Vincristine or a CENP-Ei, at concentrations 2-fold higher than the EC50 value obtained from cell viability assays in BJ Ras cells. As expected, paclitaxel significantly enhanced the mass of microtubules and vincristine reduced it. Surprisingly, the CENP-Ei significantly enhance microtubule mass (Figure 4D, Supplementary Figure 3B). MTH1i TH588 did not significantly change the whole microtubule mass, but TH1579 treatment significantly reduced the microtubule mass (Figure 4D, Supplementary Figure 3B). We next assayed microtubule dynamics by measuring microtubule plus end assembly rates in live HCT116 cells by monitoring EB3-GFP assembly. We observed a dose-dependent reduction of microtubule polymerisation rates following 1.5 hours treatment with TH588 and TH1579 (Figure 4E-F). Next, we used U2OS cells stably expressing H2B-GFP and Cherry-labelled Tubulin and found that Cherry-labelled Tubulin polymerization was impaired in a dose dependent manner of TH1579 as well as TH588 treatment (Figure 4G, Supplementary Figure 3C, 4A). Using FRAP, we observed an increased mobility of Cherry-Tubulin after 24 hours (Supplementary Figure 3D-I) and also already after a 2 hour treatment with 1 μM TH1579 (Figure 4 G-I, Supplementary Figure 3J-M) in both interphase and mitotic cells. Similar results were obtained with 5 or 10 μM TH588 (Supplementary Figure 4B-G), but Cherry-Tubulin mobility did not reach the same level as after treatment with 1 μM TH1579. It is notable that at concentrations corresponding to a cellular EC50 value i.e. 250 nM TH1579 [15], the effect upon tubulin mobility was less pronounced (Figure 4G-H) and more resembled the effects observed following 96 h MTH1 siRNA treatment in mitotic cells (Figure 3C-E). Interestingly, AZ19, albeit at relatively higher concentrations (50 μM), caused mitotic arrest and cell death (Figure 2G-H, Supplementary Figure 2B-C), and also enhanced tubulin mobility similarly to what was observed with MTH1 siRNA or TH1579 (Supplementary Figure 5 A-H). It is interesting to note that AZ19 does not show a direct effect on tubulin polymerization *in vitro* (Supplementary Figure 5I). Taken together, these data indicate that MTH1i TH588 and TH1579 appear not to substantially impact upon microtubule mass in cells, but they do significantly reduce microtubule dynamics, consistent with the mitotic phenotype, and this effect is likely owing to the direct inhibition of microtubule polymerization in combination with targeting MTH1. As shown above, we can detect interactions between MTH1 and alpha-tubulin in pulldown experiments. Therefore, by treating U2OS cells with inhibitors prior to cell lysis and pulldown of MTH1-GFP, we investigated whether cytotoxic and non-cytotoxic MTH1i could disturb this interaction. The cytotoxic (TH588 and TH1579) and non-cytotoxic MTH1i (AZ19, AZ21, IACS-4759) significantly disturbed (>70%) the interaction, while the non-cytotoxic MTH1i TH7238 and CENP-Ei showed some disturbed interaction (<40%) and the non-cytotoxic MTH1i NPD9348, as well as vincristine did not significantly disturb the interaction (Figure 4J-K).

**Fig. 4.**
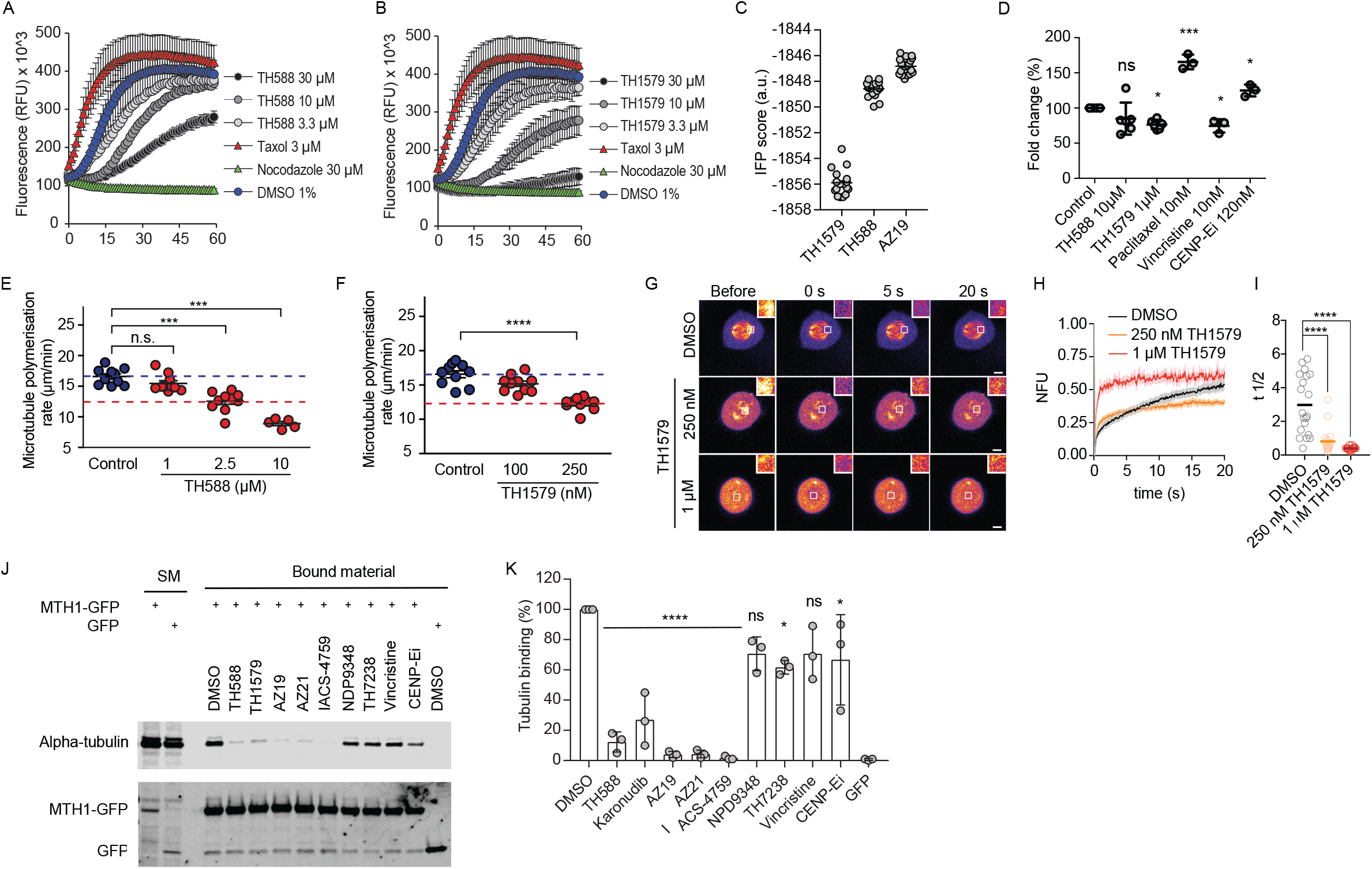
MTH1i TH588 and TH1579 reduce microtubule dynamics. **A-B** Tubulin was incubated with the indicated concentrations of MTH1i and allowed to polymerize over time, resulting in an increase in fluorescence. TH588 and TH1579 inhibited tubulin polymerization. The curves show the average result of two independent experiments. **C.** IFD scores of the induced-fit docking poses obtained by docking AZ19, TH588, and TH1579 to the colchicine binding site of tubulin. Mean values are marked by solid lines (***: p<0.001, One-way ANOVA). **D.** Quantification of whole cell microtubule analysis using flow cytometry to determine tubulin polymerization in HCT116 cells treated with 10 μM TH588, 1 μM TH1579, 10 nM paclitaxel, 10 nM vincristine or 120 nM CENP-Ei for 24 h. Data is shown as fold change of the control cells of 3 independent experiments. Statistical comparison is performed using one-way ANOVA (mean±SD, ns: non significant, *: p<0.05, ***, p<0.001). **E-F.** Measurement of mitotic microtubule plus end assembly rates, using EB3-GFP assay, in living HCT116 cells treated with the MTH1i TH588 (E) or MTH1i TH1579 (F) for 1.5 h. Scatter dot plots show average assembly rates (10 individual microtubules per cell, mean +/-SEM, t-test, n=5-10). **G-I.** Determination of microtubule dynamics in U2OS Cherry-Tubulin cells using FRAP analysis. Representative confocal FRAP image series of mitotic (G) cells treated with DMSO, 250 nM or 1 μM TH1579 for 2 h. Insets show 3 times magnified bleach area. Quantification of fluorescence recovery after photobleaching in imitotic cells (H, I). Halftime of recovery was quantified. Scatter dot blots of ≥ 20 cells from ≥ 2 independent experiments are shown. Scale bar, 5 μM. **J,K.** Tubulin binding to MTH1-GFP is reduced upon MTH1i treatment. U2OS cells expressing MTH1-GFP or GFP were treated with indicated compounds for 4 h and Alpha-tubulin bound to MTH1-GFP was detected by immunoprecipitation and Western blot. Protein levels in the starting material (SM) is shown. Values are from 3 independent experiments (mean±SD, ns: non significant, *: p<0.05, ****: p<0.0001, One-way ANOVA).

### MTH1i trigger the spindle assembly checkpoint, promoting incorporation of oxidized nucleotides into DNA and cytotoxicity

To gain additional insights into the mitotic defect caused by our MTH1i, we characterised the mitotic phenotype following treatment with these small molecules and compared the phenotype with other mitotic inhibitors. MTH1i causes a congression defect resembling the mitotic phenotype obtained following treatment of cells with a CENP-Ei, but not the microtubule poisons vincristine or paclitaxel (Figure 5A). Given this similarity we investigated potential spindle assembly checkpoint (SAC) activation by staining cells with anti-CENP-E and anti-centromere (CREST) antibodies. In control cells, the SAC is activated in prometaphase cells, as demonstrated by CENP-E and CREST co-localisation, which disappears during metaphase, when the checkpoint is inactivated (Figure 5B). Interestingly, following MTH1i treatment, CENP-E and CREST co-localisation only occurs on chromosomes misaligned from the metaphase plate, suggesting these misaligned chromosomes arrest cells in mitosis through the SAC. To test this directly, we siRNA depleted cells of MAD2 (Supplementary Figure 6A) which strongly decreased the toxicity of MTH1i (Figure 5 C-D), and CENP-Ei as previously reported [21] (Figure 5E). Furthermore, we saw a decrease in mitotic arrest using live cell imaging (Supplementary Figure 6B) and H3-S10 phosphorylation (Figure 5F), suggesting these cells survive by reducing time in mitosis. Taken together, these data indicate that cell-death following TH588 or TH1579 treatment is dependent upon a functional SAC to sustain the duration of mitotic arrest.

**Fig. 5.**
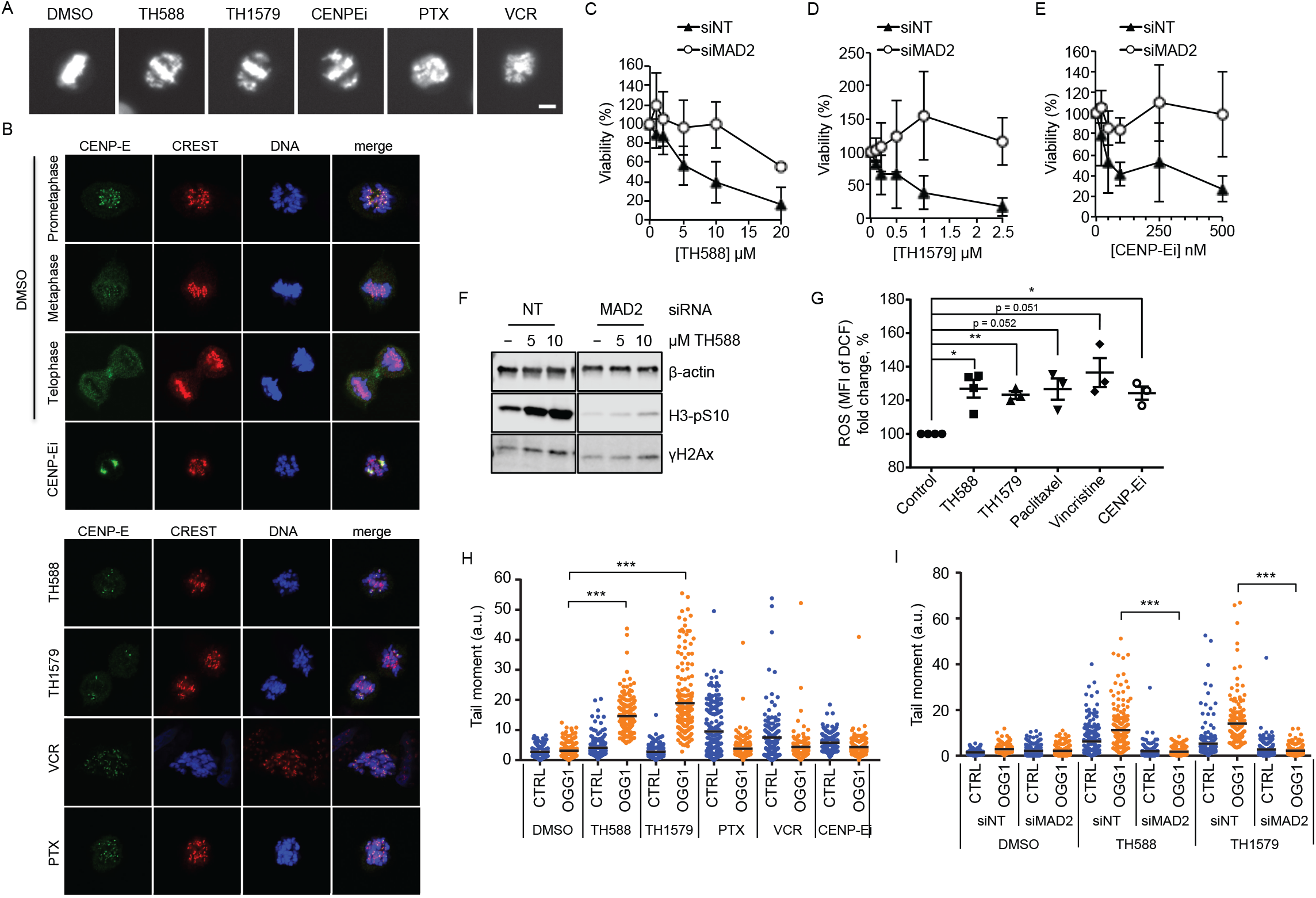
TH588 and TH1579 activate the SAC and MAD2 siRNA reduce toxicity and 8-oxodG incorporation by TH588 and TH1579. **A.** MTH1i TH588 and TH1579 show similar mitotic defect as CENP-Ei. Images of U2OS H2B-GFP cells arrested in mitosis after treatment with 5 μM TH588, 250 nM TH1579, 50 nM CENP-Ei (GSK923295), 50 nM Paclitaxel and 25 nM Vincristine. Scale bar: 10 μm. **B.** U2OS cells treated with 100 nM CENP-Ei, 5 μM TH588, 500 nM TH1579, 50 nM Vincristine or 100 nM Paclitaxel for 6 h and immunostained with CENP-E and anti-centromere (CREST) antibodies. Scale bar: 20 μm. **C-E.** MAD2 depletion reduces the toxicity of MTH1i and CENP-Ei. Resazurin viability assay of NT RNA (filled triangles) and MAD2 siRNA (open circles) transfected cells (72 h) treated with TH588, TH1579 and CENP-Ei for additional 72 h. Values were normalized to the DMSO control (0 μM) and expressed as mean ± SD from 3 independent experiments. **F.** Western blot of U2OS cells transfected with NT RNA and MAD2 siRNA for 72 h followed by treatment with TH588 for 24 h. **G.** MTH1i and other mitotic inhibitors increase ROS accumulation. ROS was determined using the fluorescence H2DCFDA/DCF ROS accumulation assay in U2OS cells treated with 10 μM TH588, 1 μM TH1579, 50 nM Paclitaxel, 200 nM Vincristine or 100 nM CENP-Ei for 24 h. Data is shown as fold change of the control cells of 3-4 independent experiments. Statistical comparison performed using Student’s t-test (*, p<0.05; **, p<0.01) **H.** Quantification of 8-oxodG levels in DNA by the modified comet assay in U2OS cells treated with 10 μM TH588, 500 nM TH1579, 100 nM Paclitaxel, 50 nM Vincristine and 100 nM CENP-E for 24 h. Samples were treated without (CTRL) or with OGG1 to measure 8-oxodG in DNA. (200 cells in 2 exp., p<0.001, One-way ANOVA). **I.** MAD2 depletion prevents the MTH1i induced incorporation of 8-oxodG into DNA. U2OS cells transfected with NT RNA and MAD2 siRNA for 72 h followed by treatment with 5 μM TH588 and 1 μM TH1579 for 24 h. The 8-oxodG levels were analyzed with the modified comet assay (200 cells in 2 experiments, p<0.001, One-way ANOVA).

There is emerging evidence connecting mitotic arrest with ROS [22], through mitophagy [23]. To test if MTH1i or mitotic inhibitors themselves may trigger ROS, we measured the generation of intracellular ROS by using a DCFDA cellular ROS assay. Treatment of cells with TH588, TH1579, as well as paclitaxel, vincristine or a CENP-Ei, increased ROS accumulation (Figure 5G, Supplementary Figure 6C-G). This is in line with mitotically arrested cells accumulating ROS [22, 23] and with our previous experiments demonstrating an increase in protein carbonylation following MTH1i treatment [6].

Next, we evaluated the relationship between oxidised nucleotides, such as 8-oxodG, present in DNA and mitotic arrest. TH588 and TH1579 were associated with 8-oxodG accumulation in DNA, but not paclitaxel, vincristine or CENP-Ei (Figure 5H, Supplementary Figure 6H). To ascertain if accumulation of 8-oxodG is a consequence of mitotic arrest after TH588 or TH1579 treatment, we determined whether MAD2 siRNA would alter 8-oxodG incorporation into DNA. We found that the incorporation of 8-oxodG into DNA is dependent on a functional SAC to induce mitotic arrest (Figure 5I, Supplementary Figure 6I). Taken together, these data demonstrate that mitotic arrest leads to increased ROS levels, and in-turn, this is required for the increase in genomic oxidised nucleotides.

### Inhibition of microtubule polymerisation dynamics synergizes with inhibition of MTH1-dependent 8-oxodGTPase activity

Our revised mechanism of action for TH588 and TH1579 suggests our compounds arrest cells in mitosis by perturbing microtubule dynamics, both directly and indirectly, resulting in an increase in ROS and incorporation of oxidised nucleotides into DNA. Previously, repair/replication synthesis has been reported to occur in mitosis [24] and here we suggest that this mitotic repair/replication synthesis is responsible for incorporation of oxidized dNTPs into DNA (Rudd et al., submitted, https://doi.org/10.1101/573931). Hence, MTH1 may play a pivotal role in preventing oxidized dNTP incorporation during mitosis, which may be lethal.

To test if incorporation of 8-oxodGTP into DNA is toxic and if MTH1 is important in preventing such lethal incorporation, we injected 8-oxodGTP into zebrafish embryos followed by treatment with TH588, or other potent MTH1i that fail to kill cancer cells [12]. Interestingly, the non-cytotoxic MTH1i are equipotent to TH588 in killing zebrafish embryos injected with 8-oxodGTP (Figure 6A-B) demonstrating MTH1i is important to prevent toxic incorporation of 8-oxodGTP. Furthermore, when injecting 8-oxodGTP into MTH1 knockout or WT zebrafish embryos, the MTH1 knockout embroys showed a higher mortality compared to the WT embroys (Figure 6C-D). These data suggest a new dual action model for TH588 and TH1579; where mitotic arrest induces elevated levels of ROS and oxidation of dNTPs, which in the absence of MTH1 are incorporated into DNA during repair synthesis in mitosis and to some extent likely also replication, resulting in cell death (Figure 6E).

**Fig. 6.**
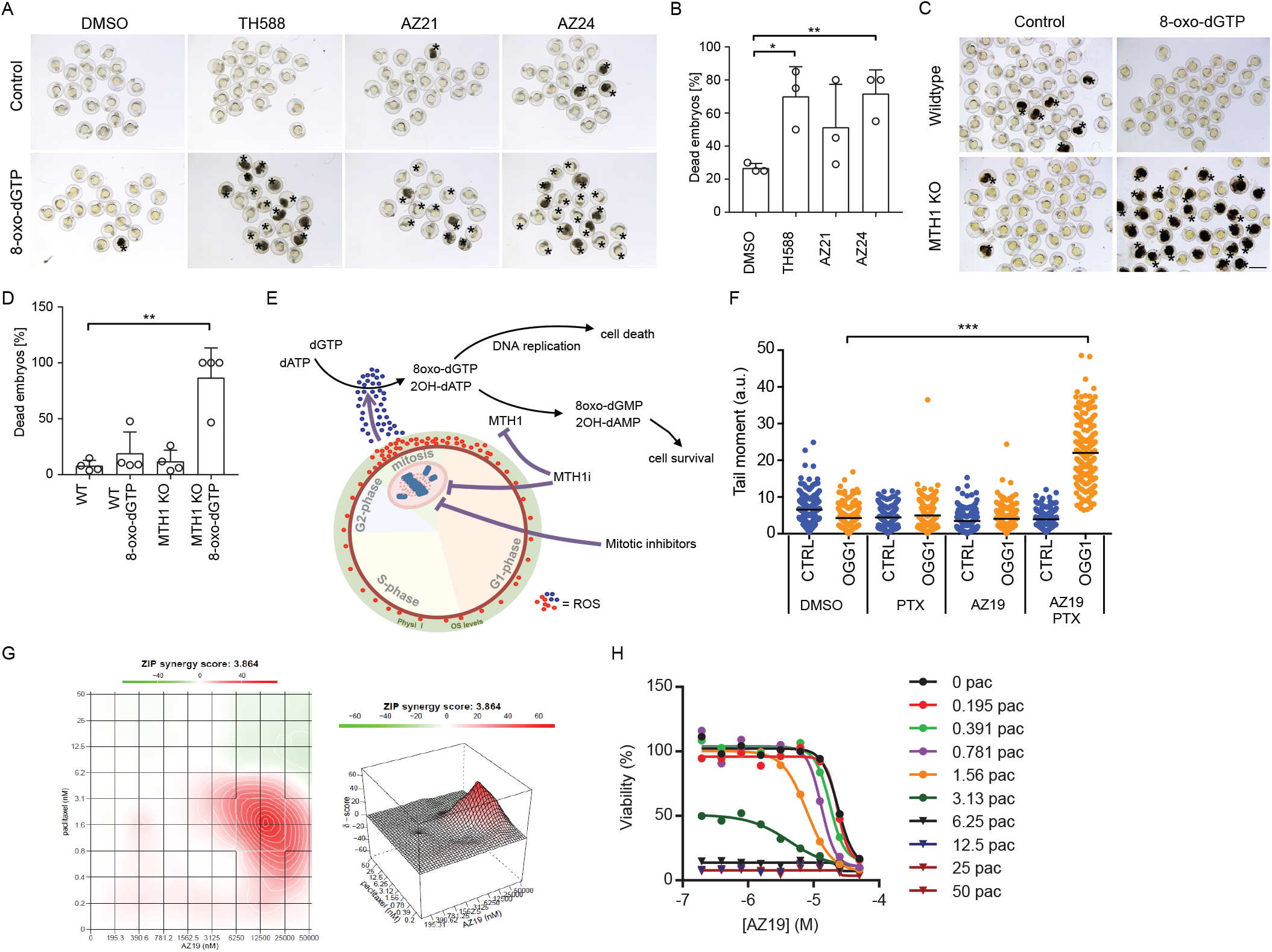
Combination of ROS accumulation by mitotic arrest and MTH1i induce 8-oxo-dG and toxicity. **A, B.** Both MTH1i causing cancer cell death or not induce zebrafish embryo lethality following addition of 8-oxo-dGTP, indicating importance of 8-oxo-dGTPase activity to induce lethality. 8-oxo-dGTP was microinjected into fertilized zebrafish embryos followed by treatment with 1.5 μM TH588, AZ21 or AZ24. **B.** 24 hours after injection, in the control group 27 ± 2 % of embryos were found to be dead whereas this number increased to 70 ± 15 % for TH588 (p = 0.014), 51 ± 21 % for AZ21 (p = n.s) and 72 ± 12 % (p = 0.01) for AZ22. Three independent experiments have been analyzed. **C, D.** MTH1 knockout embryos are sensitive to 8-oxo-dGTP injection. Wildtype or MTH1 knockout (KO) fertilized zebrafish embryos were microinjected with 8-oxo-dGTP. After 24 h, the number of dead embryos were counted, Four independent experiments were analyzed (**: p<0.01, One-way ANOVA) **E.** Schematic illustration on the proposed mechanism of action of MTH1i. Briefly, the MTH1i TH588 and TH1579 interfere with tubulin dynamics, which arrest cells in mitosis and causes an increase in reactive oxygen species (ROS), specifically in cancer cells. High levels of ROS oxidize the dNTP pool to form oxidized deoxynucleoside triphosphates, e.g., 8-oxo-dGTP and 2-OH-dATP. TH588 and TH1579 inhibit MTH1 clearance of 8-oxo-dGTP and 2-OH-dATP, resulting in incorporation of 8-oxo-dGTP and 2-OH-dATP into DNA, causing DNA damage and cell death. **F-H.** Treatment with the MTH1i, AZ19 which does not cause cancer cell death alone, leads to increased 8-oxodG incorporation into DNA and causes more cell death in combination with compounds causing mitotic arrest. **F.** Modified comet assay in A3 cells treated with 10 nM Paclitaxel and 10 μM AZ19. Quantification of comet tail moment of comets treated without (CTRL) or with OGG1 to measure 8-oxodG in DNA. (***: p<0.001, One-way ANOVA.). **G, H.** A3 cells treated with increased concentrations of Paclitaxel in combination with increased concentrations of AZ19. Viability was measured by Resazurin assay and synergy score was calculated according to the Zero Interaction Potency (ZIP) model.

A way to test this model is to separate the two mechanisms and determine if only the combined effect is synergistic in killing cancer cells. Therefore we arrested cells in mitosis with paclitaxel, to generate ROS, and treated with a non-toxic dose of AZ19. Interestingly, we observed that during paclitaxel-induced mitotic arrest the normally non-toxic AZ19 is able to introduce 8-oxodG into DNA (Figure 6F; Supplementary Figure 7A) and kill cancer cells (Figure 6G-H), similar to the effects observed with TH588 and TH1579. Similar effects were observed by arresting cells in mitosis using the CENP-Ei (Supplementary Figure 7B-C). Furthermore, combination of sub-toxic concentrations of TH1579 (125 nM) and CENP-Ei (25 nM) induced 8-oxodG incorporation (Supplementary Figure 7D-E) and combination of TH1579 with either colchicine (Supplementary Figure 8A-B), paclitaxel (Supplementary Figure 8C-D) or nocodazole (Supplementary Figure 8E-F) show synergistic effects.

## DISCUSSION

Here, we show that MTH1 has a previously uncharacterized function in mitotic progression. Following MTH1 siRNA treatment we observe a mitotic defect that is manifested by an increased time in mitosis and a decreased inter-kinetochore distance, which is a likely consequence of disrupted microtubules being unable to pull chromosomes apart. To further substantiate the role of MTH1 in mitosis, we observe MTH1 and tubulin interactions in cells, with a potential role for MTH1 to initiate microtubule nucleation.

It is established that cells accumulate ROS when arrested in mitosis [22, 23]. In line with this, we report mitotic poisons and inhibitors increases ROS, as well as TH588 and TH1579. We suggest MTH1 enzymatic activity is required under these stressed conditions to prevent incorporation of 8-oxodGTP into DNA. In support of this, no elevated levels of 8-oxodG in DNA is observed by paclitaxel, vincristine or CENPEi in spite of ROS being produced. In contrast, TH588 and TH1579 specifically elevate incorporation of 8-oxodGTP into DNA, by inhibiting MTH1-dependent 8-oxodGTPase activity. Hence, it appears that arrest in mitosis is required for ROS accumulation while concurrent MTH1i results in incorporation of 8-oxodG into DNA. We demonstrate that decreasing the time spent in mitosis using MAD2 siRNA prevented not only 8-oxodG in DNA but also cellular toxicity after TH588 or TH1579 treatment. We thus suggest that the improved survival in response to TH588 and TH1579 treatment in cells, where the spindle assembly checkpoint is disturbed, can be explained by less time spent in mitosis, ultimately resulting in lower levels of ROS generation and reduced accumulation of toxic 8-oxodG in DNA. Previously, both we and others have demonstrated that cell death caused by TH588 treatment is associated with ROS formation, supporting this hypothesis [25, 26]. Hence, MTH1i of 8-oxodGTPase only becomes relevant in presence of ROS in mitotically arrested cells.

To test if mitotic arrest and 8-oxodGTPase inhibition are both relevant for MTH1i, we combined the MTH1i AZ19 at doses that only inhibits MTH1 enzymatic activity with paclitaxel or CENPEi. The lack of 8-oxodG detection in DNA after AZ19 treatment alone is likely due to the fact that no mitotic arrest is induced with these sub-lethal doses of AZ19. Conversely, ROS induced by paclitaxel or CENPEi do not lead to 8-oxodG in DNA because of functional MTH1 activity sanitizing oxidized nucleotides. Cytotoxicity and 8-oxodG in DNA is only observed when ROS is induced by paclitaxel or CENPEi and MTH1 inhibited by AZ19. Our conclusion is that there is a dual mechanism of action to kill cancer cells, by (1) arrest in mitosis and (2) preventing sanitisation of 8-oxodGTP and other oxidized dNTPs, such as 2-OHdATP by MTH1 (Figure 6E). If the model is correct, introducing ROS by any other means than mitotic arrest would make normally non-toxic MTH1i toxic. Supporting this, we show the non-toxic MTH1i AZ21 and AZ24 are equally toxic as TH588 to zebrafish embryos upon micro-injection of 8-oxodGTP. Furthermore, we demonstrate MTH1 knockout zebrafish embryos are hypersensitive to 8-oxodGTP, demonstrating MTH1 is required for survival in presence of ROS.

Since non-toxic MTH1i kill zebrafish embryos challenged with 8-oxodGTP one would expect all MTH1i to kill cancer cells if there were high overall ROS levels in cancer. Hence, this suggests that the underlying ROS levels in cancer cells is overall too low to be toxic to cells upon MTH1i and that additional mitotic arrest to elevate ROS levels is necessary.

Cells treated with TH588, TH1579, AZ19 and also other chemically distinct MTH1i (data not shown) show similar phenotypes as observed following MTH1 siRNA treatment when it comes to congression defect, decreasing microtubule polymerisation rates, tubulin mobility, decreased inter-kinetochore distance, arrest in mitosis and generation of polynucleated cells. Importantly, the TH588 and TH1579 compounds show more pronounced mitotic arrest and effect on microtubule polymerisation as compared to AZ19, other MTH1i or MTH1 siRNA. We propose that this is likely explained by the direct effect on tubulin polymerisation shared by the TH588 and TH1579 compounds, which is not observed with AZ19. This is supported by TH588 and TH1579 being able to inhibit tubulin polymerisation *in vitro* and also affecting tubulin mobility in cells more profoundly. Since TH1579 is likely binding to the colchicine site on tubulin one would expect that the effects of TH1579 would be masked during treatments with colchicine. In contrast to this, TH1579 and colchicine potently synergize suggesting they do not share the same mechanism of action.

In conclusion, here we demonstrate that the MTH1 protein is mediating microtubule polymerisation rates required for progression of mitosis in cancer cells, which is likely by binding tubulin directly. Furthermore, some potent MTH1i break the MTH1-tubulin interaction while others do not. TH588 and TH1579 have additional effects on tubulin contributing to a stronger mitotic arrest from that observed with AZ19 or MTH1 siRNA alone. Active MTH1i work through a dual mechanism of action by inhibition of mitosis as well as inhibition of 8-oxodGTPase activity.

## AUTHOR CONTRIBUTIONS

U.W.B and T.H. conceived and supervised the project. H.G., S.R.G., N.A., L.B., L.P., K.S., C.K., A-S.J., I.A., N.S., J.B., J.M.C.M., U.WB. and T.H. designed, performed and analysed molecular pharmacological, immunostaining, biochemical and cell based experiments. E.J.H. performed computational chemistry analysis and support. L.B., C.G, N.S and U.W.B., designed, performed and analysed *in vivo* experiments. A.S., O.M., L.B., L.P., A-S.J., T.V., H. B., H.G., U.W.B and T.H designed, performed and analysed tubulin based assays. A.H., P.G and C.S., performed DNA fibre and replication stress assays. P.W., T.K., and M.S. designed and synthesised MTH1i. H.G., S.G.R., U.W.B. and T.H. wrote the paper. All authors discussed results and approved the manuscript.

