## Supplementary information for "MTH1 promotes mitotic progression to avoid oxidative DNA damage in cancer cells"

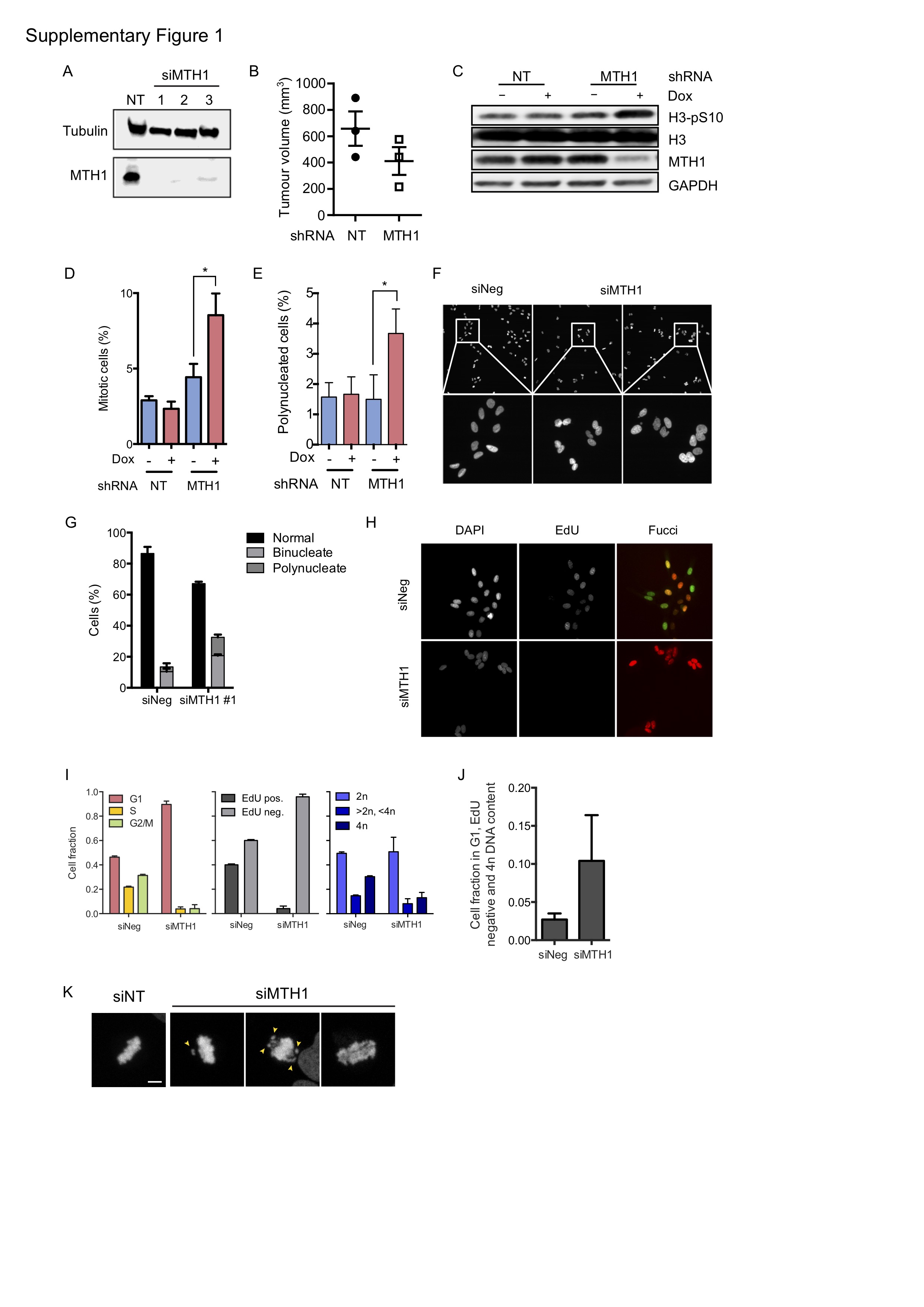

**Figure S1: MTH1 knockdown arrest cells in mitosis and induces polynucleation. A.** Western blot of U2OS cells transfected with NT RNA or MTH1 siRNA #1-3 for 10 days and stained with indicated antibodies. **B**. Tumour volume of SW480 expressing doxycycline induced shRNA against MTH1 or NT following 5 days doxycycline treatment in drinking water. Data shown as individual data and mean ± SEM. **C-E.** NT or MTH1 shRNA in HL60 cells were induced with doxycycline for 5 days and analysed by Western blot (C) or stained with Histone H3-pS10 antibodies and DAPI. H3pS10-positive cells (D) and polynucleated cells (E) were scored. **F, G**. U2OS cells were transfected with negative control (siNeg) or MTH1 (siMTH1 #1) siRNA, and 72 hours after transfection were fixed and nuclei stained with DAPI. The nuclear morphology of at least 100 cells were scored per experiment, representative images are shown. Error bars indicate SD from 3 experiments. **H-J**. U2OS-FUCCI cells were transfected with a negative control siRNA pool (siNeg) or siRNA pool targeting MTH1 (siMTH1) and 72 hours post transfection were pulsed with 10 µM EdU for 20 minutes prior to fixation and imaging to quantify cell-cycle parameters. Representative image shown in (H); proportion of FUCCI G1 phase, EdU negative, and 4n DNA content shown in (I); cell cycle distribution using FUCCI, EdU and DNA content (DAPI) in (J). Error bars indicate s.d. from single experiment. **K.** Examples of mitotic cells in U2OS cells transfected with either non-targeting (siNT) or MTH1 siRNA. Scale bar: 10 µm.

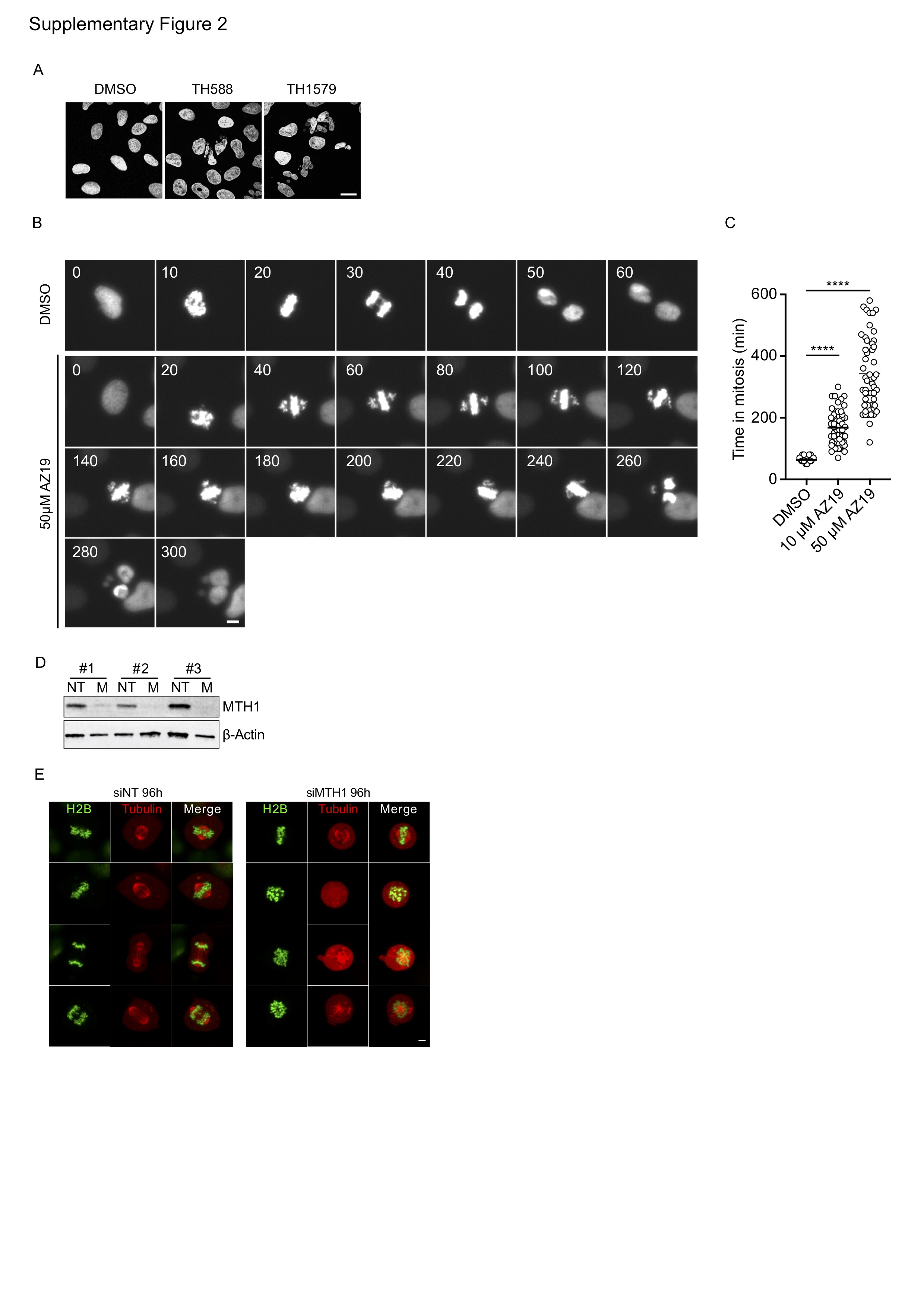

**Figure S2:** **Non-toxic AZ19 shows similar mitotic phenotype as TH588 and TH1579 at high concentrations. A.** Images of U2OS cells treated with 5 µM TH588 and 250 nM TH1579 for 24 h and stained with DAPI. **B.** U2OS H2B-GFP cells were treated with 50µM AZ19. Images were acquired in the GFP channel every 10 min. Representative images at 10 or 20 min intervals are shown. **C.** Quantification of the time in mitosis of U2OS H2B-GFP cells treated with AZ19. Values of 50 cells from 2 independent experiments are shown (****: p<0.0001, One-way ANOVA). **D.** Western blot of U2OS cells transfected with MTH1 siRNA. **E.** Representative confocal images of mitotic cells expressing H2B-GFP and Cherry-Tubulin transfected with non-targeting (siNT) or MTH1 (siMTH1) siRNA for 96 h. Scale bar, 5 µM.

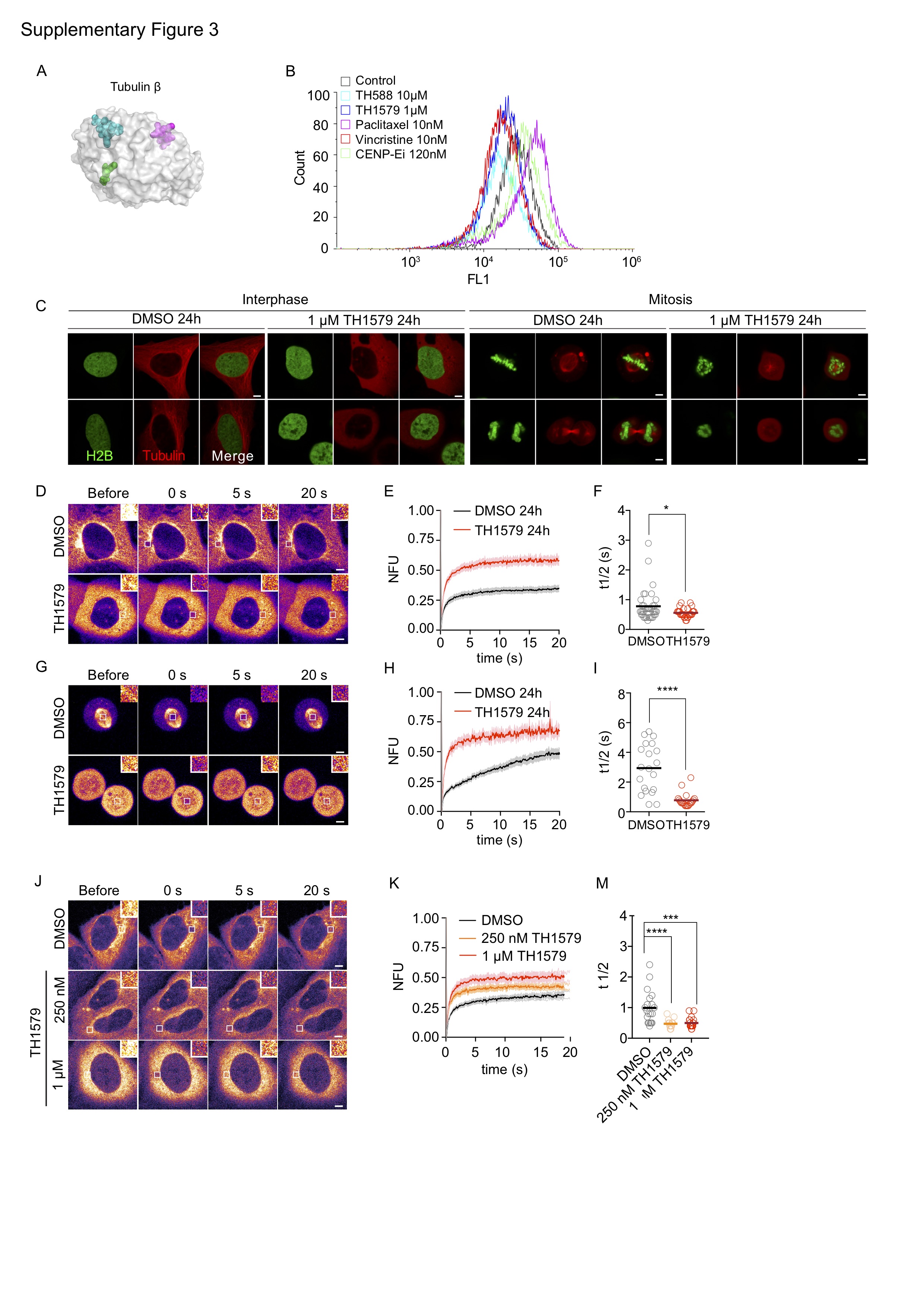

**Figure S3**: **TH588 and TH1579 destabilize microtubule formation in vitro. A.** TH588 (green spheres) docked to the colchicine binding site of β-tubulin (grey surface), showing the relative positions of bound Paclitaxel (cyan) and Vinblastine (magenta). **B.** Representative FACS plots of whole cell microtubule analysis to determine tubulin polymerization in HCT116 after 24 h treatment with indicated compounds. **C.** Representative confocal images of interphase and mitotic cells expressing H2B-GFP and Cherry-Tubulin treated with DMSO or 1 µM TH1579 for 24 h. **D, G.** Representative confocal FRAP image series of interphase (D) and mitotic cells (G) treated with DMSO or 1 µM TH1579 for 24 h. Insets show 3 times magnified bleach area. **E, F, H, I.** Quantification of fluorescence recovery after photobleaching in interphase (E, F) and mitotic cells (H, I). Halftime of recovery was quantified. Scatter dot blots of ≥ 20 cells from ≥ 2 independent experiments are shown. Scale bar, 5 µM. **J-M.** Representative confocal FRAP image series (J) and quantification of fluorescence recovery (K, M) after photobleaching in interphase cells treated with TH1579. Halftime of recovery was quantified. Scatter dot blots of ≥ 20 cells from ≥ 2 independent experiments are shown. Scale bar, 5 µM. (n.s.: non significant, **: p<0.01 ***: p<0.001 ****: p<0.0001, Student’s t-test.).

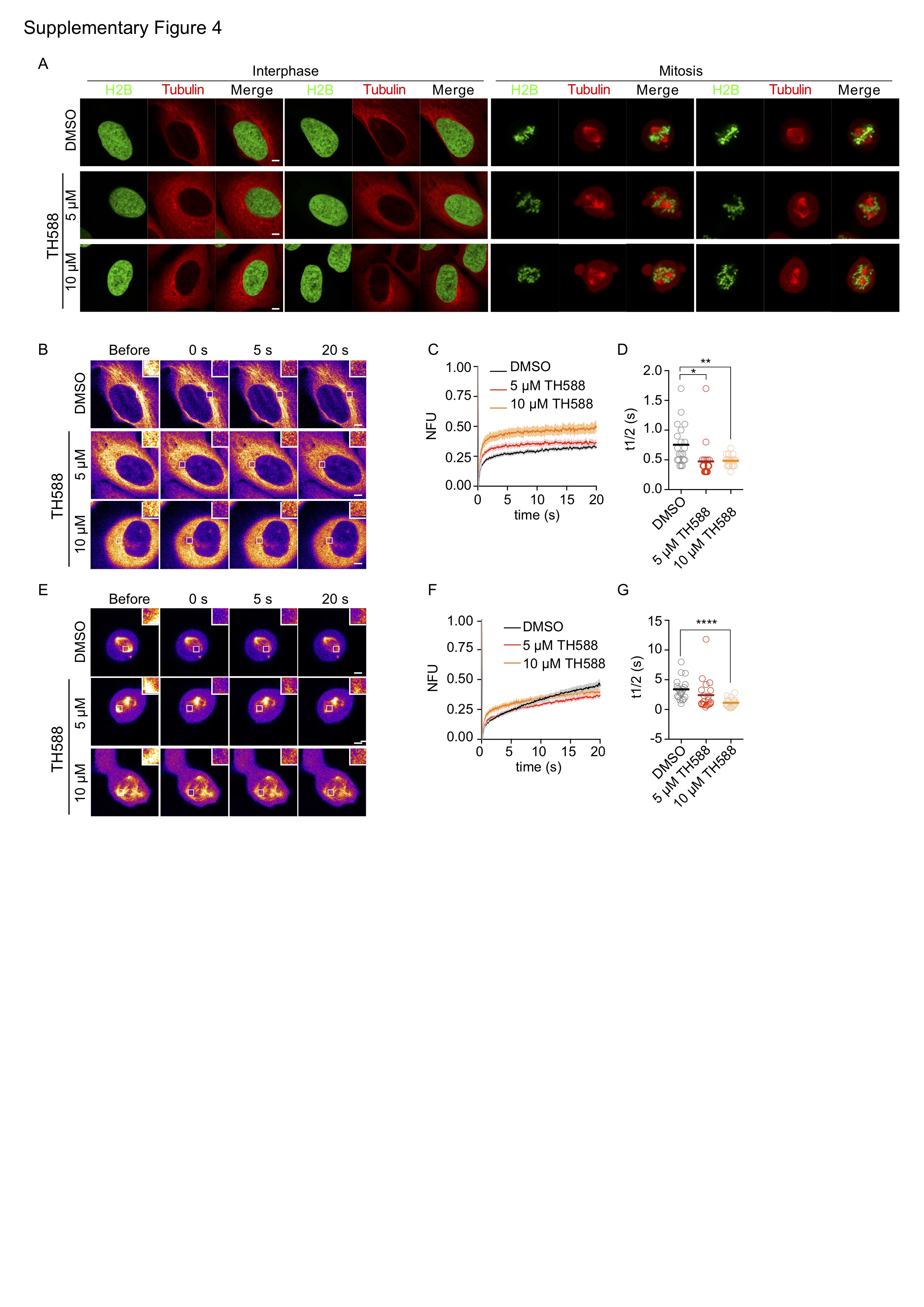

**Figure S4:** **TH588 causes microtubule disturbances. A.** Representative confocal images of interphase and mitotic cells expressing H2B-GFP and Cherry-Tubulin treated with DMSO or 5 and 10 µM TH588 for 2 h. **B, E.** Representative confocal FRAP image series of interphase (B) and mitotic cells (E) treated with DMSO or 5 and 10 µM TH588 for 2 h. Insets show 3 times magnified bleach area. Quantification of fluorescence recovery after photobleaching in interphase (C, D) and mitotic cells (F, G). Halftime of recovery was quantified. Scatter dot blots of ≥ 20 cells from ≥ 2 independent experiments are shown. Scale bar, 5 µM. (**: p<0.01 ***: p<0.001 ****: p<0.0001, Student’s t-test.)

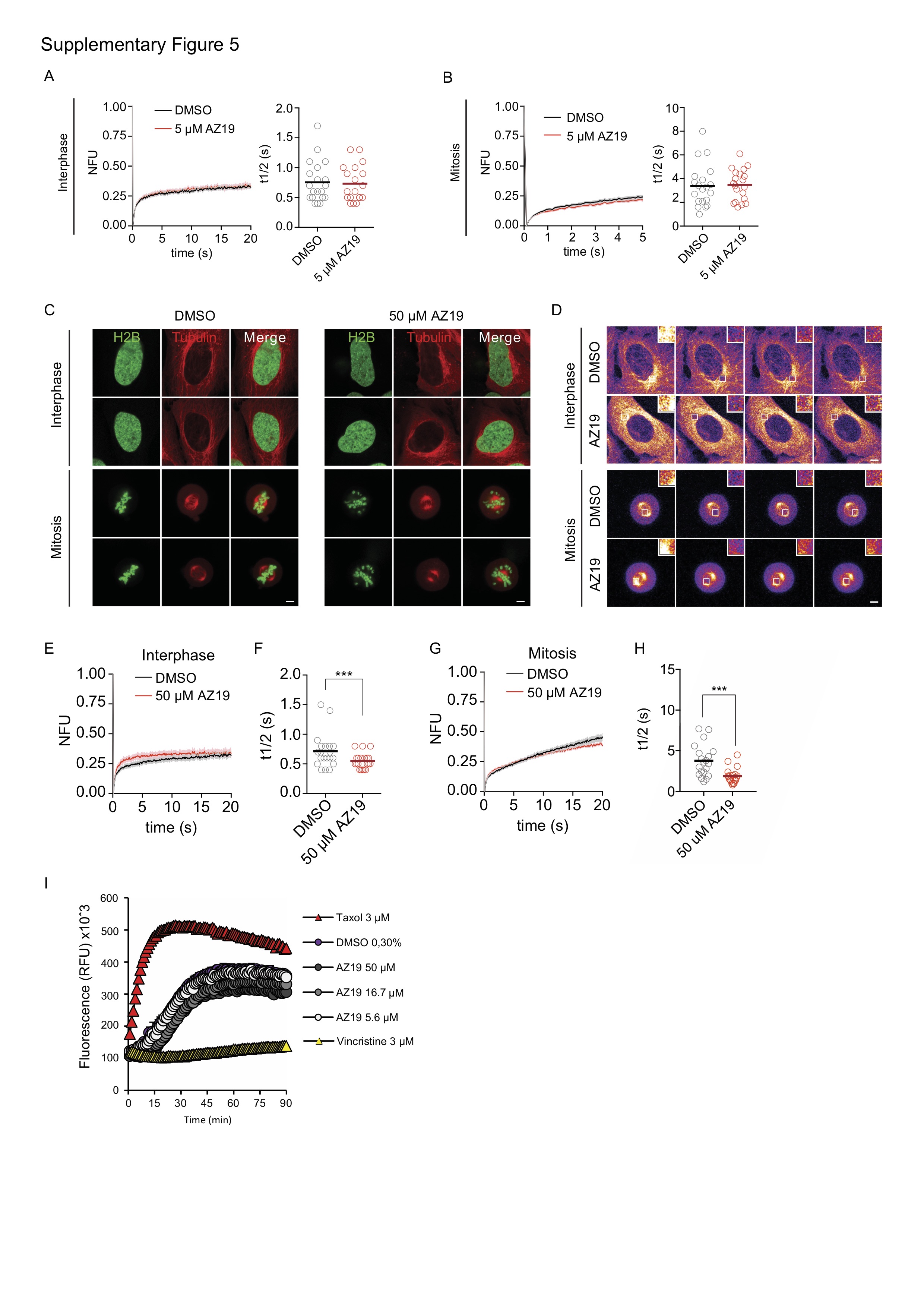

**Figure S5: Effects on microtubule organization by the MTH1i AZ19. A-B.** Analysis of microtubule dynamics after treatment with 5 µM AZ19 by FRAP analysis in interphase and mitotic cells. Quantification of fluorescence recovery after photobleaching in interphase (A) and mitotic cells (B). Halftime of recovery was quantified. Scatter dot blots of ≥ 20 cells from ≥ 2 independent experiments are shown. **C.** Representative confocal images of interphase and mitotic cells expressing H2B-GFP and Cherry-Tubulin treated with DMSO or 50 µM AZ19 for 2 h. **D.** Representative confocal FRAP image series of interphase and mitotic cells treated with DMSO or 50 µM AZ19 for 2 h. Insets show 3 times magnified bleach area. **E-H.** Quantification of fluorescence recovery after photobleaching in interphase (E, F) or mitotic cells (G, H) treated with 50 µM AZ19. Halftime of recovery was quantified. Scatter dot blots of ≥ 20 cells from ≥ 2 independent experiments are shown. Scale bar, 5 µM. **I.** In vitro tubulin polymerization assay with AZ19. Tubulin was incubated with the indicated concentrations and allowed to polymerize over time, resulting in an increase in fluorescence. The experiment was repeated twice and the curves show one representative experiment. (n.s.: non significant, **: p<0.01 ***: p<0.001 ****: p<0.0001, Student’s t-test.).

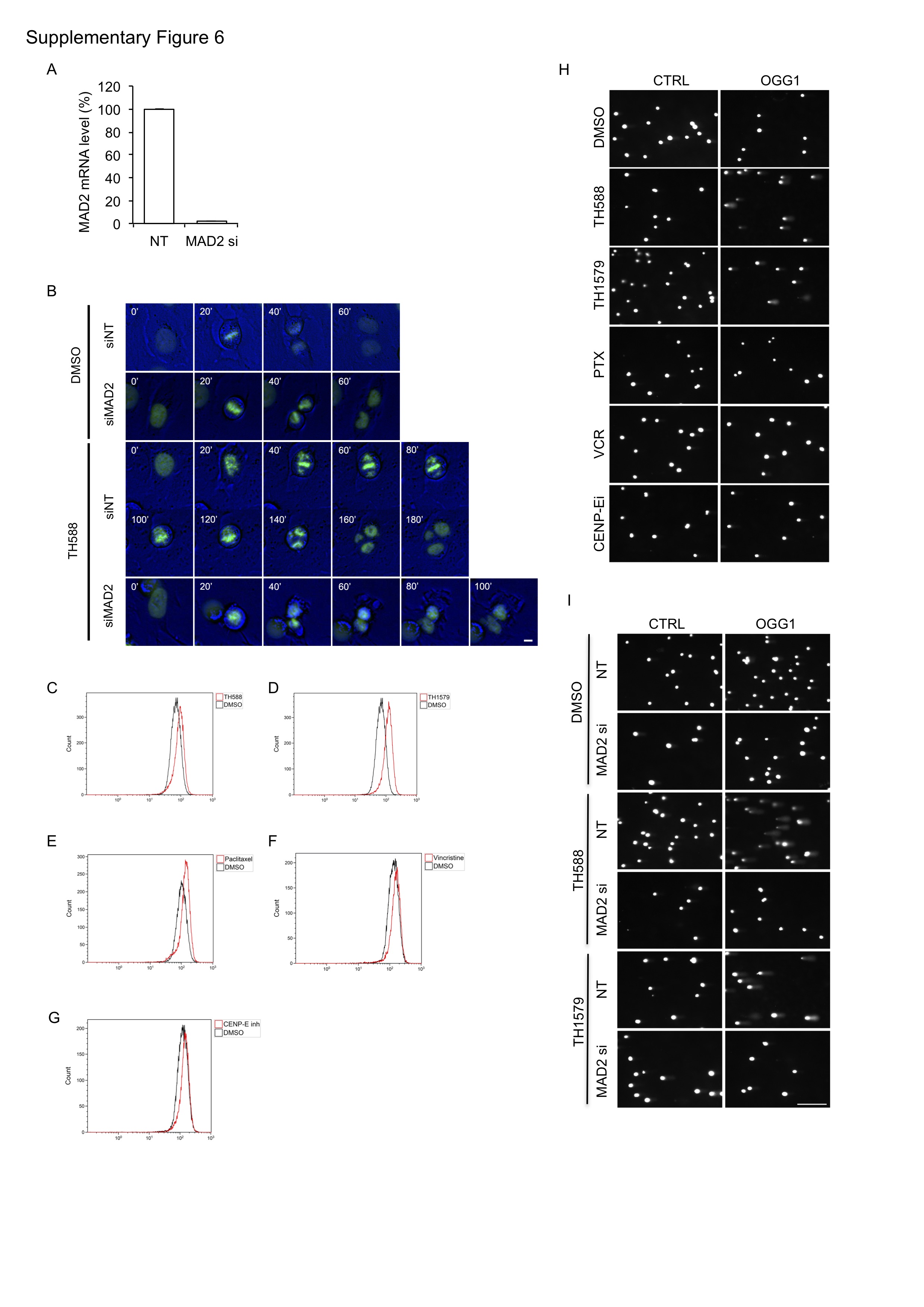

**Figure S6:** **TH588 and TH1579 induced mitotic arrest and dependent upon on SAC function. A.** mRNA levels of MAD2 measured by RT-qPCR in U2OS cells transfected with NT RNA and MAD2 siRNA for 72 h. Data shown as mean ± SD for 3 independent experiments. **B.** U2OS H2B-GFP cells were transfected with NT RNA and MAD2 siRNA for 72 h and treated with DMSO or 5 µM TH588. Images were acquired in the GFP and brightfield channels every 10 min. Representative images at 20 min intervals are shown. Scale bar: 10 µm. **C-G**. Representative FACS plots showing the fluorescent intensity of H2CDF in U2OS cells after 24 h treatment with 10 µM TH588, 1 µM TH1579, 50 nM paclitaxel, 200 nM vincristine or 100 nM CENP-Ei compared to DMSO. **H.** Representative images of comets in U2OS cells treated with 10 µM TH588, 500 nM TH1579, 100 nM paclitaxel, 50 nM vincristine and 100 nM CENP-E for 24 h. Samples were treated without (CTRL) or with OGG1. **I**. Representative images of comets of U2OS cells transfected with NT RNA or MAD2 siRNA for 72h followed by treatment with DMSO (control), 5 µM TH588 and 1 µM TH1579 for 24 h.

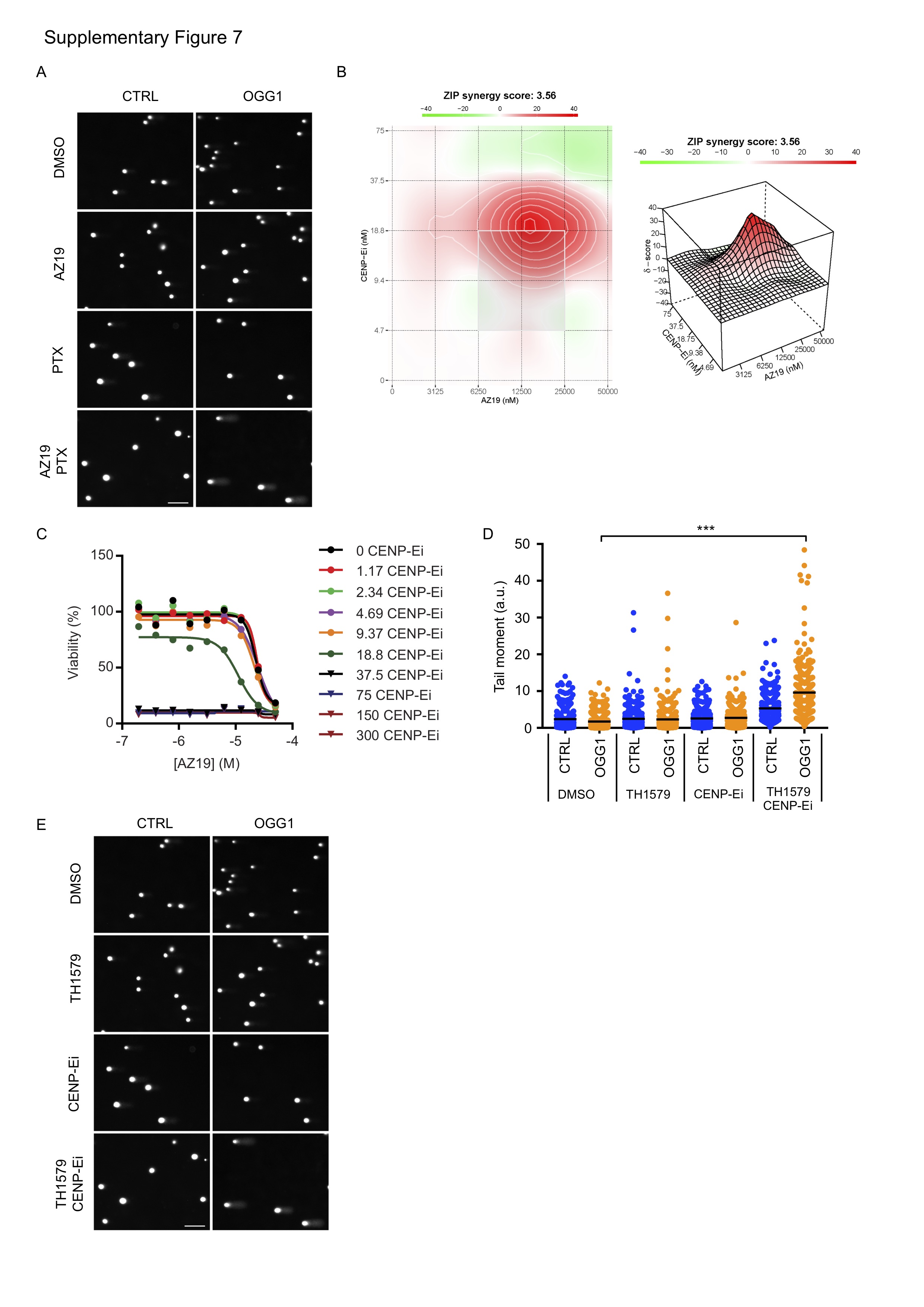

**Figure S7: Combination of a non-toxic MTH1i and mitotic inhibition induce 8-oxodG in cells.** Representative images of the modified comet assay in A3 cells treated with 10 nM Paclitaxel (PTX) and 10 µM AZ19 for 24 h. Samples were treated without (CTRL) or with OGG1 to visualize 8-oxodG lesions. **B-C**. Synergy score and viability curves of A3 cells treated with increasing concentrations of AZ19 and CENP-Ei. Viability was measured by Resazurin assay and the synergy score was calculated according to the Zero Interaction Potency (ZIP) model. **D-E.** A3 cells were treated with 125 nM TH1579 and 25 nM CENP-Ei for 24 h. The genomic 8-oxodG content was measured by the modified comet assay. **D.** Quantification of comet tail moment of comets treated without (CTRL) or with OGG1 to measure 8-oxodG in DNA. (***: p<0.001, One-way ANOVA.). **E.** Representative images of comets in U2OS cells treated with 125 nM TH1579 and 25 nM CENP-Ei.

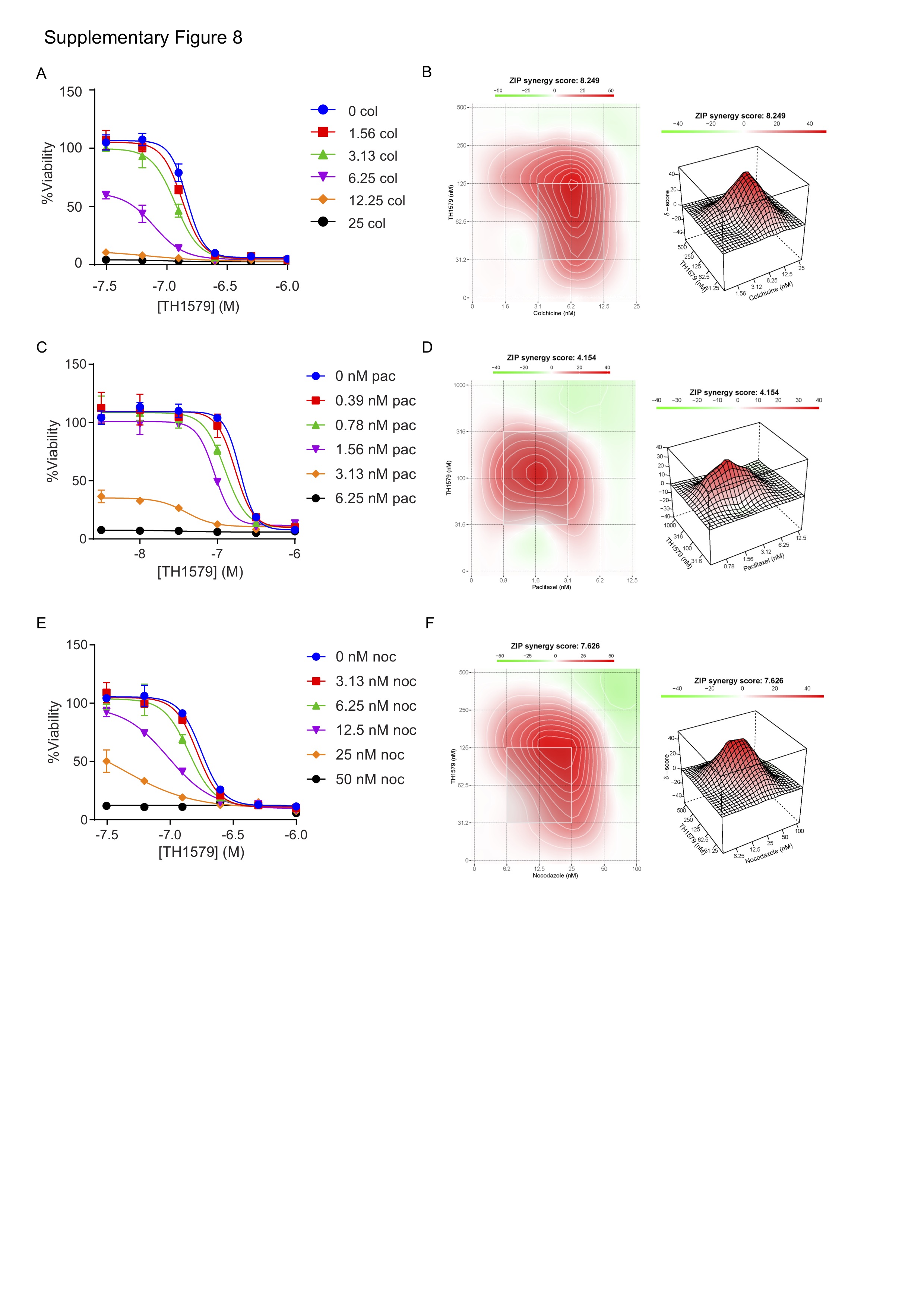

**Figure S8: Synergy between TH1579 and microtubule inhibitors.** Synergy score and viability curves of A3 cells treated with increasing concentrations of TH1579 and Colchicine (A, B), Paclitaxel (C, D) or Nocadozole (E, F)). Viability was measured by Resazurin assay and the synergy score was calculated according to the Zero Interaction Potency (ZIP) model.

**SUPPLEMENTAL MOVIES**

**Movie S1**. Time-lapse of U2OS H2b-GFP cells treated with DMSO (control) for 24h.

**Movie S2.** Time-lapse of U2OS H2b-GFP cells treated with 10 µM TH588 for 24h.

**Movie S3.** Time-lapse of U2OS H2b-GFP cells treated with 500 nM TH1579 for 24h.

**Movie S4.** HCT116 cells treated with 2 µM DME for 1.5 hrs before EB3-GFP tracking.

**Movie S5.** U2OS H2bGFP cells treated with DMSO (control) for 24h.

**Movie S6.** U2OS H2bGFP cells treated with 50 µM AZ19 for 24h.

Materials and Methods

**Cell culture**

All cell lines were cultured at 37°C in 5% CO2 and in media supplemented with 10% FBS (except A2780 cells) and Penicillin Streptomycin (100 U/ml). U2OS (ATCC) and SW480 (ATCC) cells were cultured in DMEM, A3 and HL60 (ATCC) cells in RPMI-1640 Glutamax, BJ-hTERT and BJ-hTERT/SV40T/RasV12 cells in DMEM Glutamax with 4.5 g/L glucose, A2780 Pol δ replacement cells [1] in DMEM with 5% low TET FBS and 100 µg/mL G418, 10 µg/mL Blasticidin and 1 µg/mL Puromycin. All cell culture reagents were from Gibco/Thermo Fisher. BJ-hTERT and BJ-hTERT/SV40T/RasV12 cells were a gift from W. Hahn, Harvard Medical School, Boston, MA.

**Inhibitors**

Following chemical inhibitors were used: CENP-E inhibitor (GSK923295, SelleckChem), Vincristine sulfate (Sigma Aldrich), Paclitaxel (Sigma Aldrich), Nocadozole (Sigma Aldrich), RO3306 (SelleckChem), Hydroxyurea (Sigma Aldrich), Aphidicolin (Sigma Aldrich). TH588[2], TH1579 [3], TH7238 and AZ cmpd #19, #21, #24 [4] were synthesized in house and prepared as previously reported.

**Antibodies**

Following antibodies were used: mouse anti beta-Actin (ab6276, Abcam) order number is missing), mouse anti-ATM pS1981 (sc47739, Santa Cruz Biotech), rabbit anti-ATR phospho S428 (ab178407, Abcam), mouse anti-BubR1 (NB100-353, Novus Biologicals), rabbit anti-Cdk2 phospho-T14/T15 (Santa Cruz, sc28435-R), mouse anti-CENP-E (#14977, Cell Signaling), rabbit anti-Chk1 phospho S345 (#2341, Cell Signaling), mouse anti-Chk1 (#2360, Cell Signaling), human anti-CREST (15-235-001, Antibodies Inc.), rabbit anti-GAPDH (sc-25778, Santa Cruz),mouse anti-H2A.X phospho S139 (05-636, Millipore), mouse anti-Histone H3 phospho-S10 (H3-pS10; ab14955, Abcam), rabbit anti-Histone H3 (#4318, Cell Signaling), rabbit anti-MTH1 (NB100-109, Novus Biologicals), rabbit anti-p21 (H164, Santa Cruz Biotech), rabbit anti-cleaved PARP (#9541, Cell Signaling), mouse anti-PLK1 (Millipore, #5844), mouse anti alpha-Tubulin.

**Transfection of siRNA, shRNA and plasmids for overexpression**

For the siRNA transfection, 10 nM of siRNA was used together with INTERferin transfection reagent according to the manufacturer’s instructions. Following siRNA sequences were used: MAD2 si: 5′-GGAAGAGUCGGGACCACAGUU-3′, MTH1 si#1 5’-GACGACAGCUACUGGUUUC-3’, MTH1 si#2 5’-GAAAUUCCACGGGUACUUC-3’, MTH1 si#3 5’-CGACGACAGCUACUGGUUU-3’. For non-targeting RNA, the All-Stars negative control (Qiagen) was used.

The shRNA Ginseng lentivirus vectors (Gad et al 2014) were transfected into SW480 cells and transfected cells were selected with 4 µg/ml puromycin. Following sequences were used: NT shRNA: 5’-GGAACTAGCATACGTAAGTAA-3’; MTH1 sh#2: 5’-CGAGTTCTCCTGGGCATGAAA-3’.

shRNA vectors as described in [5] were transfected into HL60 cells and transfected cells were selected with 4 µg/ml puromycin. Following sequences were used: NT shRNA: 5’-GGAACTAGCATACGTAAGTAA-3’; MTH1 sh#2: 5’-CGAGTTCTCCTGGGCATGAAA-3’.

The H2B-GFP vector were constructed by amplifying H2B from the H2B-RFP pENTR1A vector (Addgene # 22525) by PCR and subcloning the product into the pENTR1A-GFP-N2 vector (Addgene # 9364) at the HindIII and BamH1 restriction sites. The H2B-pENTR1A-GFP-N2 vector was verified by sequencing and transferred into the pLenti-CMV-Blast and pLenti-CMV-Hygro vectors using LR clonase (Invitrogen).

mCherry-Tubulin-6 was a gift from Michael Davidson (Addgene plasmid # 55147), H2B-RFP in pENTR1A (w507-1) and pENTR1A-GFP-N2 (FR1) were gifts from Eric Campeau and Paul Kaufman (Addgene plasmids # 22525 and # 19364).

H2B-GFP plasmids were transfected using lentivirus infections and mCherry-Tubulin-6 plasmid was transfected using PEI reagent.

NT or MTH1 shRNA in HL60 cells were induced with 0.5 µg/mL doxycycline for 5 days. At the end of incubation, cells were recollected for Western blot or immunofluorescence.

**Immunofluorescence**

Immunofluorescence microscopy with suspension cells was performed as described previously [6]. Mitotic cells were detected using anti-H3-pS10 antibody and nuclei were stained with 4,6 diamidino-2-phenylindole (DAPI). Approximately 1 million cells/sample were recollected for quantitative immunofluorescence analysis of H3-pS10. Cells were washed with 5% bovine serum albumin (BSA) in PBS and incubated for 5 min with 4% paraformaldehyde (PFA) in PBS at room temperature (RT). Then cells were washed again with 5% BSA in PBS.  Cells were permeabilized and blocked with 5% BSA, 0.1% Tween-20 and 0.1% Saponin in PBS. Cells were incubated for two hours at RT with rabbit anti-H3-pS10 antibody. Cells were then washed and incubated with anti-rabbit IgG Alexa Fluor 568 (1:1000; Life technologies A11036) for one hour at RT. 4,6 diamidino-2-phenylindole (DAPI) was used to stain DNA. Next, cytospin centrifuge was used to spin the cells onto microscope slides. Finally, images were acquired in a Zeiss LSM-780 confocal microscope at 63-fold magnification. More than 120 cells per sample were counted manually and cells with high H3-pS10 signal and chromatin condensation were scored as mitotic cells.

**RT-qPCR**

Cells were collected by centrifugation and resuspended in PBS. Total RNA was prepared with the Direct-zol RNA miniprep kit (Zymo Research) and cDNA was prepared with QuantiTect Reverse Transcriptase kit (Qiagen) according to manufacturer’s instructions. Each qPCR reaction was made in triplicates and the Mad2 values were normalized to the control genes ~~Beta~~ beta-Actin, GAPDH and HPRT. Each reaction contained 10 ng of cDNA, 10 µM forward and reverse primers and 1x iTaq universal SYBR green supermix (Bio-Rad). The qPCR reactions were performed in a Rotor-Gene Q instrument (Qiagen). Following primers were used: Mad2: 5’-CAGGATGAAATCCGTTCAGTG-3’, 5’-ATAAATCAGCAGATCAAATGAACAA-3’; beta-Actin: 5’-CCTGGCACCCAGCACAAT-3’, 5’-GGGCCGGACTCGTCATACT-3’; GAPDH: 5’-AAGGTCGGAGTCAACGGATT-3’, 5’-CTCCTGGAAGATGGTGATGG-3’; 5’-HPRT: 5’-GACCAGTCAACAGGGGACAT-3’, 5’-AACACTTCGTGGGGTCCTTTTC-3’.

**Western blot**

Cells were washed with 1x PBS and scraped in lysis buffer (10 mM Hepes pH 7.1, 50 mM NaCl, 0.3 M sucrose, 0.1 mM EDTA, 0.5% Triton X100, 1 mM DTT, 1x protease inhibitor cocktail (Thermo), and 1x Halts phosphatase inhibitor cocktail (Thermo)). Samples were incubated on ice for 30 min before preparation in Laemmli Sample Buffer (Bio-Rad) containing 355 mM final concentration 2-mercaptoethanol, and samples were denatured at 95°C for 5 min. Samples were loaded on 4-12% SDS-PAGE gel (Mini-Protean TGX Precast gel, Bio-Rad) and the proteins were transferred to nitrocellulose membranes using the Trans-Blot Turbo instrument (Bio-Rad). Membranes were stained with Ponceau S and blocked in 5% Milk/1%BSA in TBS/0.1% Tween for 1 h. Antibodies were diluted in TBS/0.1% Tween and IRDye 800CW and IRDye 680LT secondary antibodies were used. Images of blots were obtained using the LI-COR Odyssey Fc Imaging system.

For protein analysis of HL60 cells, approximately 1.2 million cells per condition were recollected, washed once ~~in~~ with PBS and then lysed by resuspension in Lysis buffer (150 mM NaCl, 50 mM Tris-HCl pH 7.4, 1% Triton X-100, 5 mM EDTA, 1 mM EGTA, complete EDTA-free protease inhibitor cocktail (Roche) and HALT phosphatase inhibitors (Thermo). The suspension was left on ice for 30 minutes and then sonicated for complete lysis. Next, the suspension was spun down at 12000 g at 4ºC for 15 minutes. The supernatant was recollected into a new tube. Protein concentration was determined using the Bradford assay (Thermo). 20 μg of total protein/sample was mixed with Laemmli buffer (0.4% Bromophenol blue, 8% SDS, 40% glycerol, 200 mM Tris-HCl pH 6.8 and 40 mM DTT), warmed at 95ºC for 10 min and loaded on 4–20% precast polyacrylamide gel (Mini-PROTEAN® TGX™ Precast Gels, Bio-Rad) for proteins separation. After that, the proteins were transferred to nitrocellulose membrane. Thereafter membrane was blocked in Odyssey Blocking Buffer (Li-Cor) for 1 h at RT and incubated overnight with primary antibodies at 4ºC. antibodies for 1 h at RT. The secondary antibodies used were: anti-Mouse IgG-IRDye 800CW (Li-Cor) and anti-rabbit IRDye 680RD (Li-Cor). Membrane was visualized using Odyssey Fc Imaging System (Li-Cor). Next, membrane was washed and stripped with Restore PLUS Western Blot Stripping Buffer (Thermo) for 15 minutes.

**Fluorescence recovery after photobleaching (FRAP)**

U2OS cells stably expressing GFP-tagged H2B and Cherry-tagged Tubulin were incubated in phenol-red free medium containing 0.1% DMSO or TH1579 (1µM or 250 nM) for indicated time periods. Cells were transferred to a Zeiss LSM780 confocal microscope equipped with a UV-transmitting Plan-Apochromat 40x/1.30 Oil DIC M27 objective. eGFP and mCherry were excited with a 488 nm Ar laser and a 561 nm DPPS laser, respectively. The microscope was equipped with a heated environmental chamber set to 37°C.

For FRAP analysis, a 40x40 pixel sized ROI in the cytoplasm was selected using the Regions tool of the ZEN software (ZEN, Zeiss, Germany) and photobleached with the 561 nm laser set to maximum power at 100% transmission using 5 iterations at scan speed 8 (5 µs). Before and after bleaching, confocal image series were recorded with the following settings: 100 ms time intervals (20 prebleach and 200 postbleach frames), frame size 256x256 pixels, 110 nm pixel size, bidirectional scanning and a pinhole setting of 2.52 airy units. Mean fluorescence intensities of the bleached region were corrected for background and for total nuclear loss of fluorescence over time. For the quantitative evaluation of photobleaching experiments, data of at least 18-36 nuclei from two-three independent experiments were averaged and the mean curve, the standard error of the mean (s.e.m.), halftime of recovery was calculated and displayed using Microsoft Excel software and GraphPad Prism. Statistical significance was calculated using Students t-test.

**F3H assay**

To analyse the interaction of GFP-tagged MTH1 with alpha-Tubulin in cells, we employed the previously described F3H assay [7, 8]. The U2OS F3H cell line was purchased from ChromoTek (Chromtek, Germany) and transfected with a GFP-MTH1 expression vector. After 24-48 h, cells were fixed in 4% PFA permeabilized in 0.5% Triton X100 and probed with an antibody against alpha-Tubulin (Abcam, ab18251) for 1 h at RT. After three washes in PBST, secondary antibody donkey-anti-rabbit Alexa Fluor 647 (Life Technologies, A21245) was added for 1 h at RT together with DAPI. Cells were imaged on a ZEISS LSM780 with a Plan-Apochromat 40x/1.3 Oil DIC M27 objective and sequential scanning.

**High content microscopy**

Treated cells, either 48 hours post siRNA for U2OS or 3 days with 10 ng/ml doxycycline for Pol δ replacement cells, were seeded at 10 000 cells per well in black, clear-bottomed 96-well plates (BD Falcon). The following day, if labeling S-phase cells, cells were pulse-labelled with 10 µM EdU for 20 minutes prior to fixation in 4% paraformaldehyde in PBS (Santa Cruz) containing 0.5% Triton X-100 for 15 min. Wells were washed with PBS before permeabilisation in 0.3% Triton X-100. For detection of EdU labelled cells, a click reaction mix was assembled in PBS containing 2.5 mM copper (II) sulphate, 10 mM ascorbic acid and 2 µM ATTO-488 azide fluorophore (ATTO-TEC), which was added to wells and incubated in the dark for 30 minutes. Wells were washed in PBS, cell nuclei stained with 1 µg/ml DAPI for 10 min, and further washed in PBS prior to imaging. Plates were imaged on an ImageXpress high-throughput microscope (Molecular Devices) with a 20x objective. Images were analyzed (nuclei counting, DAPI and EdU nuclear intensity measurements) with CellProfiler (Broad Institute) and data handled in Excel and plotted in Prism6.

**Modified comet assay**

Cells were seed in 6-well plates at a density of 150-200 000 cells per well and either on the same day (suspension cells) or the day after (adherent cells) treated for 24h with compound or DMSO control . Cells were harvested by trypsinization, washed once in PBS and resuspended in 300µl PBS. The suspension (100 µl) was mixed with 500 µl 1.2% low melting point agarose at 37°C and the mixture were added to agarose coated slides and a coverslip was added on top. The slides were solidified on ice and lysed O/N at 4°C in Lysis buffer (2.5 M NaCl, 100 mM EDTA, 10 mM Tris, 10% DMSO, 1% Triton X100). Samples were washed 3 times in Enzyme buffer (40 mM HEPES, 0.1 M KCl, 0.5 mM EDTA, 0.2 mg/ml BSA, pH 8.0) and treated with hOGG1 enzyme (2 µg/ml) or buffer alone for 45 min at 37°C. hOGG1 was expressed in *E. coli* and purified as previously described (Gad et al 2014). Slides were transferred to Alkaline Electrophoresis buffer (300 mM NaOH, 10 mM EDTA) for 30 min and electrophoresis was performed at 25V, 300 mA for 30 min at 4°C. Samples were washed in 400 mM Tris pH 7.5 for 45 min. DNA was stained with SybrGOLD and comets were quantified with Comet Assay IV software in live video mode.

**Time-lapse microscopy**

Cells were seeded in 96-well plates (BD Falcon plate 353376; 5000 cells/well). The day after, cells were treated and the time-lapse was initiated after 30 min and images of GFP and brightfield channels were acquired every 10 min for 24 h in a Perkin-Elmer ImageXpress instrument. Cells were kept at 37°C in 5% CO_2_ atmosphere during the entire time-lapse experiment. Movie files were assembled in MetaXpress and ImageJ softwares. For each condition, images were acquired from two different wells. Individual cells were followed manually and scored for defects in mitosis including mitotic slippage/polynucleation (MS/PN), micronuclei formation (G1/MN), mitotic slippage (MS) or cell death during mitosis (DiM). The time in mitosis was defined as the time from the cell rounding up and condensing the chromatin to the end of cytokinesis.

***In vitro* tubulin polymerization**

Tubulin polymerization was measured *in vitro* using a fluorescence-based tubulin polymerization assay kit (BK011P from Cytoskeleton Inc.) with modifications. Briefly, 2 µg/µl tubulin was incubated in the supplied reaction buffer using a final concentration of 15% glycerol and 1 % total concentration of DMSO in a final volume of 20 µl in black 384-well plates (Optiplate, Perkin-Elmer). Fluorescence was recorded as function of time at 37°C in a Hidex sense plate reader using 355(40) and 436(20) filters for excitation and emission, respectively.

**Measurement of microtubule plus end assembly rates by EB3-GFP**
Microtubule plus end growth rates were determined by tracking EB3-GFP protein in living HCT116 cells. Cells were transfected with pEGFP-EB3 (provided by Linda Wordeman, USA) and siRNAs for MTH1 as indicated, seeded onto glass bottom dishes (Ibidi, Germany) and after 48 hours cells were treated with the Eg5/Kif11 inhibitor Dimethylenastron (2 μM, Sigma, USA) in the presence or absence of MTH1i, paclitaxel, NAC and KBrO3, respectively as indicated. Four sections with a Z-optical spacing of 0.4 μm were taken with an Olympus 60X 1.42 NA objective every two seconds using a Deltavision ELITE™  microscope equipped with a PCO Edge® sCMOS camera (PCO, Germany) at 37°C, 5% CO2. Images were deconvolved using the SoftWorx 6.0 software (Applied Precision, USA). Average microtubule assembly rates (μm/min) were calculated based on data received for 20 individual microtubules per cell and a total of 10–30 cells were analyzed.

**Whole cell microtubuli analysis by flow cytometry**

HCT116 cells were seeded in 6 well plates the day before, treated with the different compounds as indicated for 24 hours, and finally cells were harvested by trypsination and subjected to the staining protocol for whole cell microtubule[9] analysis as previously described [9]. Cells were analyzed by flow cytometry (Navios, Beckman Coulter) using the FL1 channel.

### **Detection of intracellular ROS by flow cytometry**

The generation of intracellular ROS was determined using the fluorescent probe 2’,7’-dichlorodihydrofluorescein diacetate (H2DCFDA; Molecular Probes). The non-fluorescent H2DCFDA passively diffuses into cells and is converted to the highly fluorescent 2’,7’-dichlorofluorescein (DCF) upon oxidation by ROS.

U2OS cells were seeded in 6 well plates and exposed to compounds 24 hours after. Following an exposure of 12 hours, cells were harvested by trypsination and resuspended in serum-free medium. After adding H2DCFDA to a final concentration of 10 µM, cells were incubated for 30 min at 37°C and analyzed by flow cytometry (Navios, Beckman Coulter) using the FL1 channel.

**Zebrafish**

Zebrafish were raised and staged according to standard protocols. For injection experiments, 8-oxo-dGTP (Trilinkbiotech #2746), was mixed with injection buffer (9 µM spermine, 0.21 µM spermidine and 0.3 % phenol red) to a final concentration of 1.3 mM. 1-2 nl were injected into 1-cell stage zebrafish embryos. Directly after, embryos were transferred to a 6-well plate (20 embryos/well) and exposed to chemicals as indicated. 24 hours after injection, embryos were scored and disintegration or severe morphological abnormalities were regarded as dead. All experiments were performed in accordance to the local ethical guidelines.

The zebrafish tupfel-longfin (TL) wildtype line as well as the MTH1 knock-out line Nudt1^UU1732^ line was used. The latter was generated using the CrispR/Cas9 technique by the Genome Engineering Zebrafish facility, Scilifelab, Uppsala, Sweden. Homozygous loss of MTH1 was confirmed by genotyping.

**Xenograft mouse study**

The animal study was performed in compliance with EU 2010/63 directive and approved by the regional experimental animal ethical committee in Stockholm (N484/12). The animals were housed under sterile conditions and were provided with enrichment and free access to food and water. The environmental conditions, temperature, humidity, cage size and light cycle were according to laboratory animal guidelines and regulations.

SW480 cells transfected with MTH1 or non-targeting (NT) shRNA #2 expressing doxycycline inducible vector were subcutaneously inoculated in the flank of 5 weeks old, female SCID mice (locally bred at Karolinska Institute, Sweden). Tumours were measured by a caliper and calculated as width x length x depth. When tumours reached a volume of approximately 250 mm^3^, 2 mg/ml doxycycline was added to the drinking water. Five days following doxycycline treatment, three animals inoculated with either SW480 MTH1 shRNA or SW480 NT shRNA expressing vectors were euthanized, tumours dissected and placed in 4% PFA for at least 24 hours before change to 70% ethanol.

**Mitotic aberration quantified in xenografts**

Xenografts from mice with or without MTH1 shRNA suppression were collected, paraformaldehyde fixated and paraffin embedded at day four after start of doxycycline treatment. The Paraffin embedded xenografts were sectioned with a thickness of 5 μm and stained with hematoxylin and eosin. All cells in metaphase and anaphase on each xenograft section were counted and classified as normal or aberrant. Only cells with fully developed metaphase plates were considered as metaphases. Metaphases were defined as aberrant if they had a spitted metaphase plate. Anaphases were defined as aberrant if they displayed bridges, lagging chromosomes, chromosomes stuck at the poles or were multipolar.

**In silico docking studies**

All molecular modeling studies were performed using Schrödinger Suite 2016-02 (Schrödinger LLC, New York, NY). Images were prepared using Pymol 1.8.2 (Schrödinger LLC, New York, NY).

Protein preparation: the tubulin-colchicine complex published by Prota *et al.* (4O2B.pdb),[10] was used for docking. The structure was prepared with the Protein Preparation Wizard. After preprocessing, only chain A (α-tubulin with bound GTP) and chain B (β-tubulin with bound colchicine) were retained. Protonation and metal charge states were generated for the hetero groups, and the most likely states were retained. The hydrogen bonding network was then refined by sampling water orientations and optimizing the states for hydroxyls, Asn, Gln, and His residues. After that, waters with less than 3 H-bonds to non-waters were removed. Finally the complex was minimized with restrained heavy atoms sand unrestrained hydrogens using the OPLS3 force field, until a heavy atom convergence of 0.3 Å was reached.

Ligand preparation: the structure of TH588 was manually sketched and then converted to a low-energy 3D conformation using LigPrep. Epik was used to predict different tautomers and protonation states. The state with the lowest energy was then used for docking.

Induced-Fit docking: in order to allow for structural adaptation to TH588, protein side-chain flexibility was simulated using the standard Induced-Fit Docking protocol as implemented in the Schrödinger software [10, 11]. The bound colchicine was used to define the center of the docking grids. Glide SP was used for both docking stages, and the OPLS3 force field was used for minimizations.
